# Post-metamorphic skeletal growth in the sea urchin *Paracentrotus lividus* and implications for body plan evolution

**DOI:** 10.1101/2020.10.09.332957

**Authors:** Jeffrey R. Thompson, Periklis Paganos, Giovanna Benvenuto, Maria Ina Arnone, Paola Oliveri

## Abstract

Understanding the molecular and cellular processes that underpin animal development are crucial for understanding the diversity of body plans found on the planet today. Because of their abundance in the fossil record, and tractability as a model system in the lab, skeletons provide an ideal experimental model to understand the origins of animal diversity. We herein use molecular and cellular markers to understand the growth and development of the juvenile sea urchin (echinoid) skeleton. We developed a detailed staging scheme based off of the first ∼four weeks of post-metamorphic life of the regular echinoid *Paracentrotus lividus*. We paired this scheme with immunohistochemical staining for neuronal, muscular, and skeletal tissues, and fluorescent assays of skeletal growth and cell proliferation to understand the molecular and cellular mechanisms underlying skeletal growth and development of the sea urchin body plan. Our experiments highlight the role of skeletogenic proteins in accretionary skeletal growth and cell proliferation in the addition of new metameric tissues. Furthermore, our work provides a framework for understanding the developmental evolution of sea urchin body plans on macroevolutionary timescales.

## 1. Introduction

The evolution of animal body plans has resulted in the vast diversity of animal morphologies seen in deep time and on the planet today [1]. Crucial for understanding the evolution of animal morphology is, however, a precise understanding of the molecular and cellular mechanisms which operate during animal growth and development [2]. Skeletons, the hard, biomineralized tissues which provide structure and support for numerous animals, make up the majority of the animal fossil record, but are also ideal systems for understanding the genetic and cellular changes that take place during body plan development and evolution [3]. Echinoderms, the clade of deuterostomes including starfish and sea urchins, have a biomineralized endoskeleton made up of a porous meshwork of CaCO_3_ and comprising numerous interlocking and abutting skeletal plates [4]. The echinoderm skeleton has conferred upon the group an exceptional fossil record, precisely demonstrating evolutionary changes in body plans [5, 6]. Within the echinoderms, the sea urchin (echinoid) larval skeleton has also become a paradigm for understanding the mechanistic basis for development, and the genetic regulatory networks operating in larval echinoid skeletal development are exceptionally well-understood [7, 8]. While there is a breadth of knowledge concerning the development and evolution of the larval skeleton in sea urchins, there remains a poorer understanding of the development of the juvenile sea urchin skeleton at the level of the gene, protein, and cell.

Most echinoids, like many marine invertebrates, have a biphasic life style. Following a protracted larval stage, the adult or juvenile body plan emerges from the larvae during metamorphosis [9]. The post-metamorphic sea urchin body plan is pentaradially symmetrical, globe shaped, and comprised of multiple CaCO_3_ tessellate plates which make up the test (Fig. 1). Depending upon the species, test plates are covered in one or multiple protrusions called tubercles, which attach to spines via a ball-and-socket joint (Fig. 1). The skeletal tissues of the echinoid test have been historically demarcated into axial and extraxial structures [10]. Axial structures include the ocular and ambulacral plates, skeletal plates through which the tube feet of the water vascular system protrude, and the interambulacral plates, which, in juveniles, are characterized by large primary spines (Fig. 1a,b). New axial skeletal elements are added to the test from a growth zone at the margin of the ocular plates (Fig. 1a) [11]. Extraxial elements of the echinoid test are confined to the most aboral skeletal elements of the test, and include the periproctal and genital plates (Fig. 1a). Extraxial elements contrast with axial elements because they are not added via a distinct growth zone.

**Figure 1.**
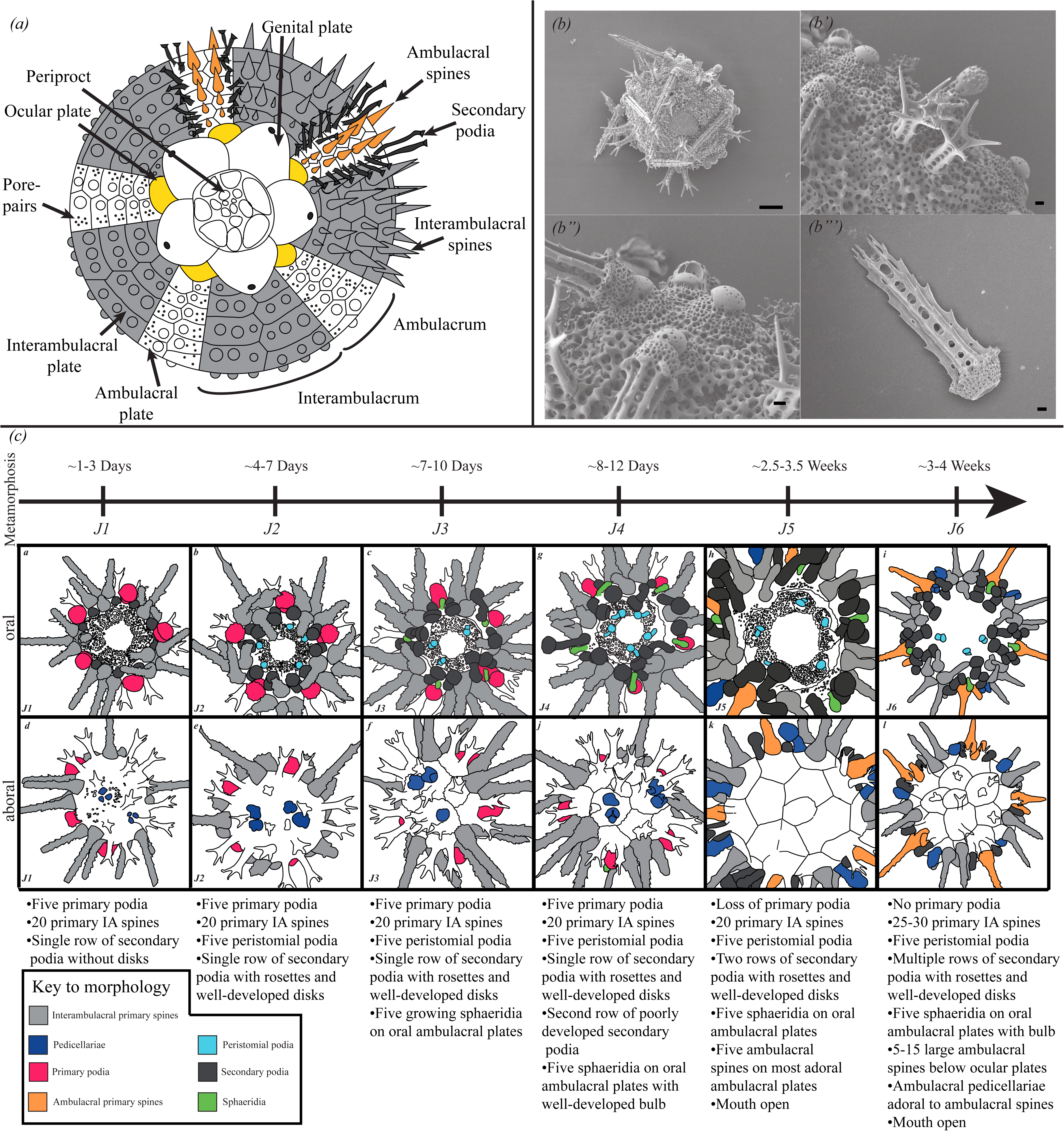
(a) Diagram modified from [43] showing morphology of adult sea urchin test from adoral view (b) SEM image showing aboral view of test with juvenile and primary spines Scale bar is 100 μm. (b’) Close up of b showing juvenile spines, ambulacral spine, and sphaeridia (b’’) Close up of b showing the details of the interambulacra. Primary interambulacral spines articulate with the bosses of primary tubercles on interambulacral plates. (b’’) Isolated primary spine. Scale bars in b’-b’’’ 10 μm. (c) Staging scheme showing example drawings and descriptions of each stage and the addition of morphological structures through the first 4 weeks of development. Structures are color coded as in the figure legend.

Growth of the echinoid test has been attributed to two distinct processes, plate addition and plate accretion [12]. Plate addition is characterized by the formation of new test plates at the margin of the ocular plates [11]. As new plates are added, and the animal grows larger, the position of these plates relative to the ocular plate shifts closer to the oral surface. This process is tightly linked to the addition of other metameric structures of the echinoid test such as the secondary podia [13-15]. Plate accretion is the accumulation of new skeletal material onto the margins of already-added plates. This process results in the elaboration of test plate morphology, and is reminiscent of growth and remodeling of vertebrate bone [16].

Despite the long history of work characterizing changes in echinoid morphology through ontogeny [9, 11, 12, 17-21], there has been relatively little work tying these morphological changes to their molecular and cellular underpinning. To fill this gap, we performed detailed analyses of morphological changes during growth, immunohistochemistry, and fluorescent assays of skeletal growth and cell proliferation in post-metamorphic juveniles of the sea urchin *Paracentrotus lividus*. This has shed light on the molecular and cellular processes operating during juvenile sea urchin growth, with a focus on the processes of skeletal plate addition and accretion.

## 2. Material and Methods

### (a) Culturing and staging scheme

Two cultures of larvae from different pairs of adult *P. lividus* were reared through to metamorphosis at the culturing facilities at the Stazione Zoologica Anton Dohrn (Supplemental Methods). After metamorphosis, which took place over the period of about two to three days, approximately fifty post-metamorphic juveniles were present in each culture. As cultures developed, animals were examined daily using a ZEISS Stemi 2000 stereomicroscope, imaged at different times post-metamorphosis and scored for the presence or absence of morphological structures to establish a staging scheme.

### (b) Immunohistochemistry and imaging

Juvenile and late stage *P. lividus* larvae were fixed following protocol in [22] and described in Supplemental Methods. Primary and secondary antibodies used are found in Supplemental Table 1. Specimens were imaged using a Zeiss LSM 700 confocal microscope, Leica SPEinv inverted confocal microscope using sequential scanning, or Zeiss Lightsheet Z1 microscope.

### (c) Assays of skeletal growth and cell proliferation

To visualize newly deposited CaC0_3_ of the skeleton, live animals were incubated with Calcein (Sigma) as described in [23]. To understand the spatial distribution and quantity of proliferating cells during juvenile sea urchin growth assays of cell proliferation were carried out using the Click-iT® EdU Alexa Flour® 555HCS (Life Technologies) and Click-iT™ EdU Cell Proliferation Kit for Imaging Alexa Flour™ 647 (Thermo Fisher Scientific) following [23]. Cells were quantified for analyzed images using ImageJ and R. Additional details of calcein staining, proliferative staining and imaging can be found in Supplemental Methods.

## 3. Results

### (a) Stages of juvenile growth

During growth of the late larvae, the adult body plan develops (Supplemental Fig. 1) and the pentaradial juvenile emerges during metamorphosis [9]. After metamorphosis, juveniles are approximately 200-400 μM in diameter and have distinct ambulacral and interambulacral areas with spines and primary podia. Juveniles grow rapidly following metamorphosis, adding additional structures such as spine-like sensory sphaeridia and pincer-like pedicellariae (Fig. 1b,c). Post-metamorphic growth is asynchronous across and within cultures. Thus to make standardized comparisons across different individuals, morphological characteristics of ∼100 animals from two cultures were analyzed daily from 0 to 4 weeks post-metamorphosis and grouped based on morphological differences. This resulted in a staging scheme based on the presence or absence of morphological features encompassing six distinct stages herein termed J1-J6 (Fig. 1c; Supplemental Fig. 2). Stages are summarized in Fig. 1, and a detailed description is given in the Supplemental Data.

**Figure 2.**
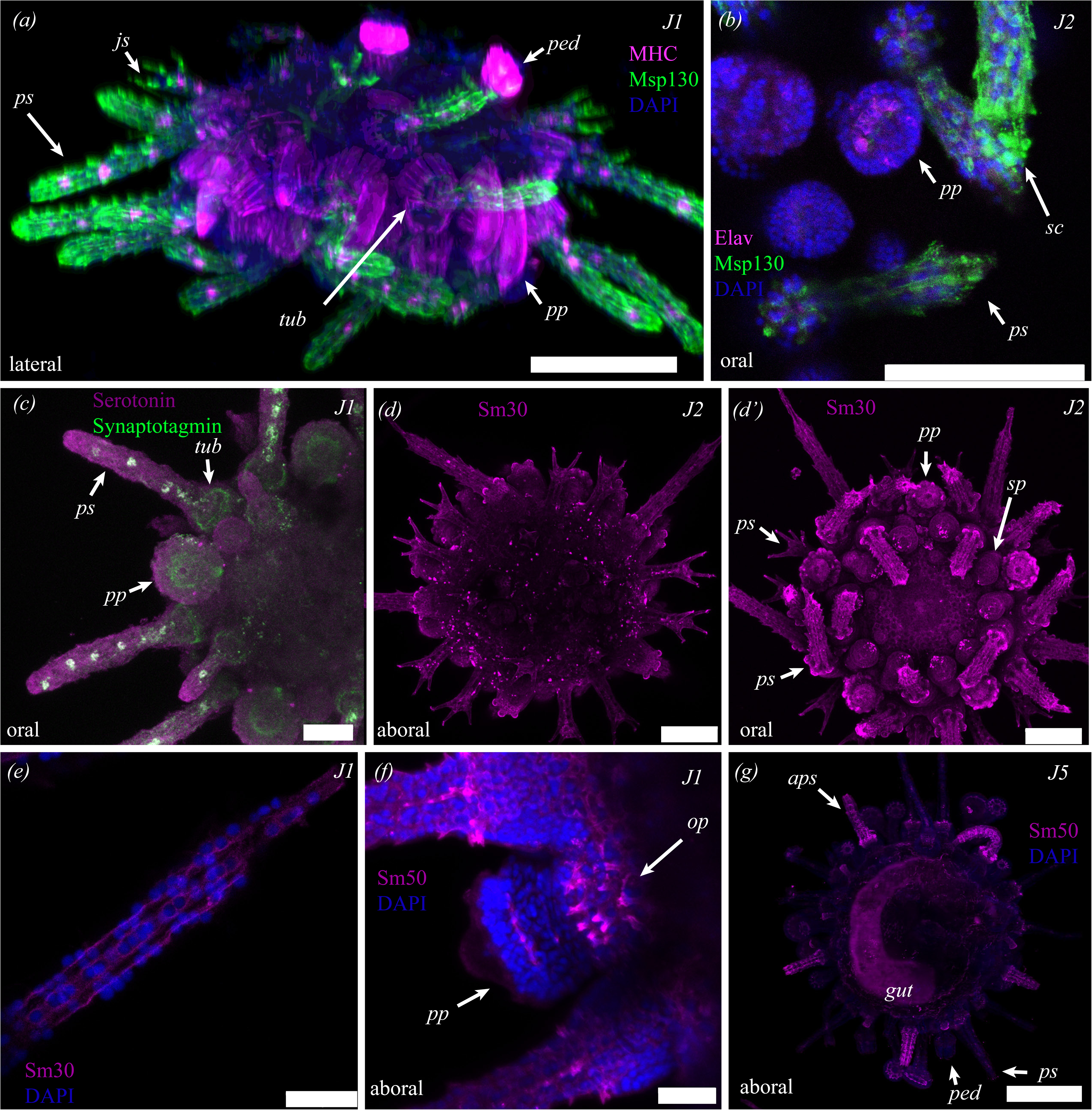
Molecular characterization of juvenile *P. lividus* cell and tissue types. (a) Staining with antibodies against MSP130 and Myosin Heavy Chain reveals the distribution of musculature. Details in text. (b). Staining against ELAV and MSP130 shows nerves in the secondary podia, and skeletogenic cells in the spines. (c) Staining for synaptotagmin and serotonin reveals the extent of the nervous system and serotonergic neurons. Details in main text. (d-g) Immunostaining using antibodies against the skeletogenic proteins SM30 and SM50 stains skeletal tissues, with stronger staining in more recently deposited biomineral. js, juvenile spine; ps primary interambulacral spine; ped, pedicallariae; aps, ambulacral primary spine; pp, primary podia; sp, secondary podia; tub, tubercle; gut, gut; op, ocular plate, gp, genital plate; ap, anal plate; sph, sphaeridia; al, Aristotle’s lantern. Scale bars in a, b,d, d’ 100 μm, c 50 μm, e-f 25μm, and g 200 μm.

### (b) Molecular characterization of juvenile tissues

In order to understand the molecular basis for the development of the structures identified in our staging scheme, immunohistochemical markers were used to characterize tissue and cell types in *P. lividus* juveniles (Fig 2a-g). To visualize muscular tissues, an anti-Myosin Heavy Chain (MHC) antibody that identifies and localizes muscles surrounding the gut of larval echinoderms [24] was used. Staining in a J1 individual (n=1) with the anti-MHC antibody shows reactivity along the interior of the tube feet (Fig. 2a, Supplemental Fig. 3), in longitudinal bands surrounding the base of spines and tubercles (Fig. 2a, Supplemental Fig. 3), in bundles along the length of the spines (2a, Supplemental Fig. 3) and, most strongly in the base of the pedicellariae (Fig. 2a, Supplemental Fig. 3). This is comparable to immunoreactivity recently shown for F-actin [25].

**Figure 3.**
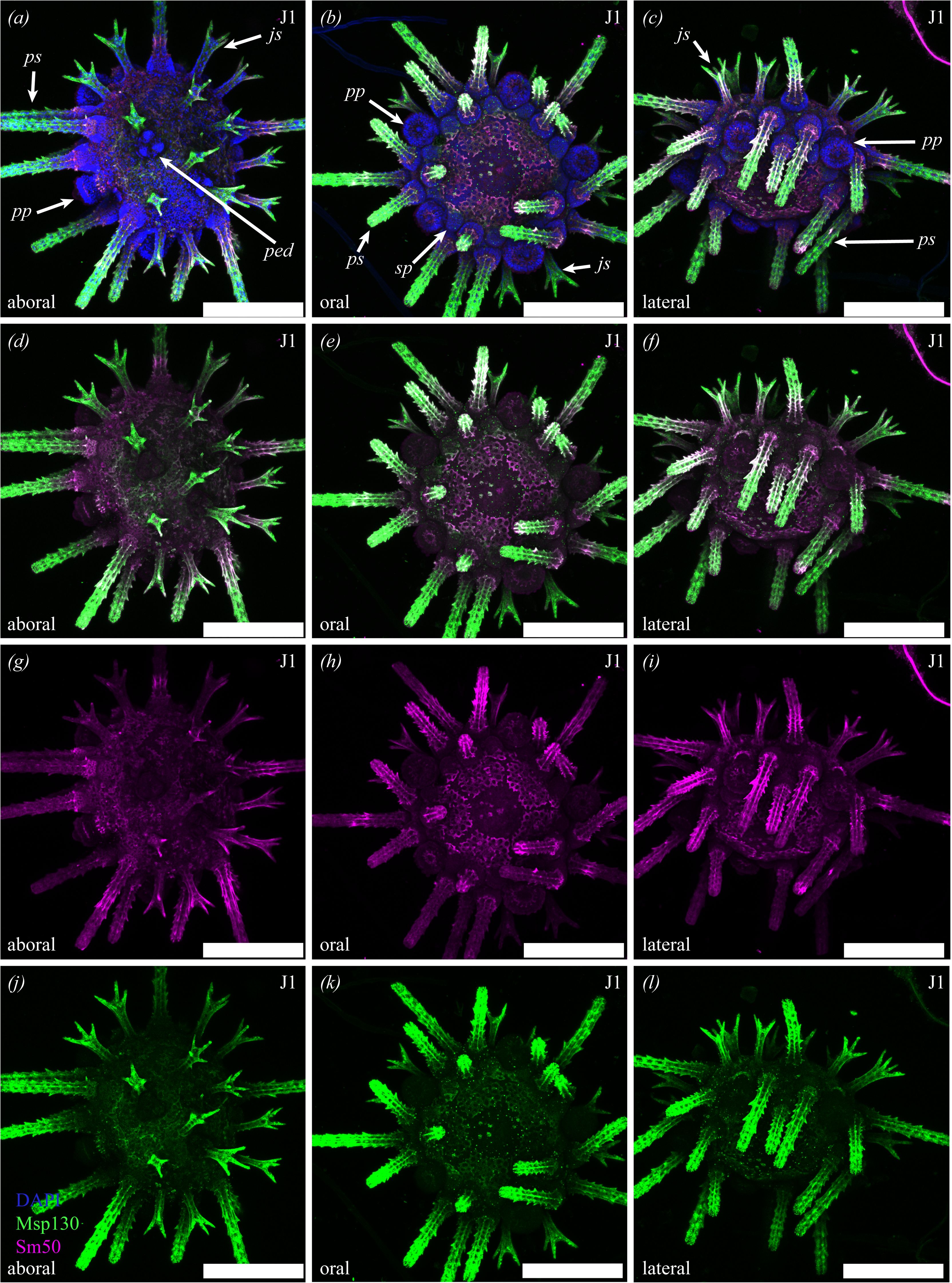
Localization of the skeletogenic proteins MSP130 (green) and Sm50 (purple) in a J1 *P. lividus* juvenile. Description of staining given in the main text. Abbreviations as in Figure 2. Scale bars 200 μm.

To identify and characterize the structure of the juvenile nervous system, samples were incubated with antibodies against pan- and specific neuronal markers. These results are largely in agreement with a recent characterization of the *P. lividus* nervous system. [25] 1E11, a pan-neuronal marker of the synaptic vesicle trafficking protein synaptotagmin [26], was used to identify the distribution of neurons, neuronal projections, and synapses (Fig. 2c, Supplemental Fig. 4). 1E11 identified bundles of fibers that form a ring around the bases of the tubercles, in the tube feet, and in cells along the lumen of the spines (n=5;Fig. 2c). In contrast to the pan-neuronal synaptotagmin, an antibody against serotonin identified a smaller subpopulation of serotonergic neurons along the lumen of the interambulacral primary spines where they co-localize with synaptotagmin. (n=1;Fig. 2c, Supplemental Fig. 4). Another neuronal marker is ELAV, an RNA-binding protein used to identify neurons [26]. Isolated ELAV^+^ cells were found in the primary and secondary podia (n=1;Fig. 2b, Supplemental Fig. 5) in a pattern reminiscent of sensory motor neurons found in podia of ophiuroids [27]. Importantly, these cells are distinct from the skeletal tissues of the rosette (Fig. 2b, Supplemental Fig. 5). Co-staining with anti-ELAV and 1E11 revealed co-reactivity in cells in the primary interambulacral spines. (Supplemental Fig. 6). Acetylated tubulin has also been used to characterize neurons in other animal groups. An anti acetylated-tubulin antibody reacted with numerous cilia covering the surface of J1 individuals, and distributed in a circlular pattern around the margins of the distal ends of podia. Staining is also present along the interior of newly forming secondary podia, which may be the neural plexis of [25] (n=2;Supplemental Fig. 7).

**Figure 4.**
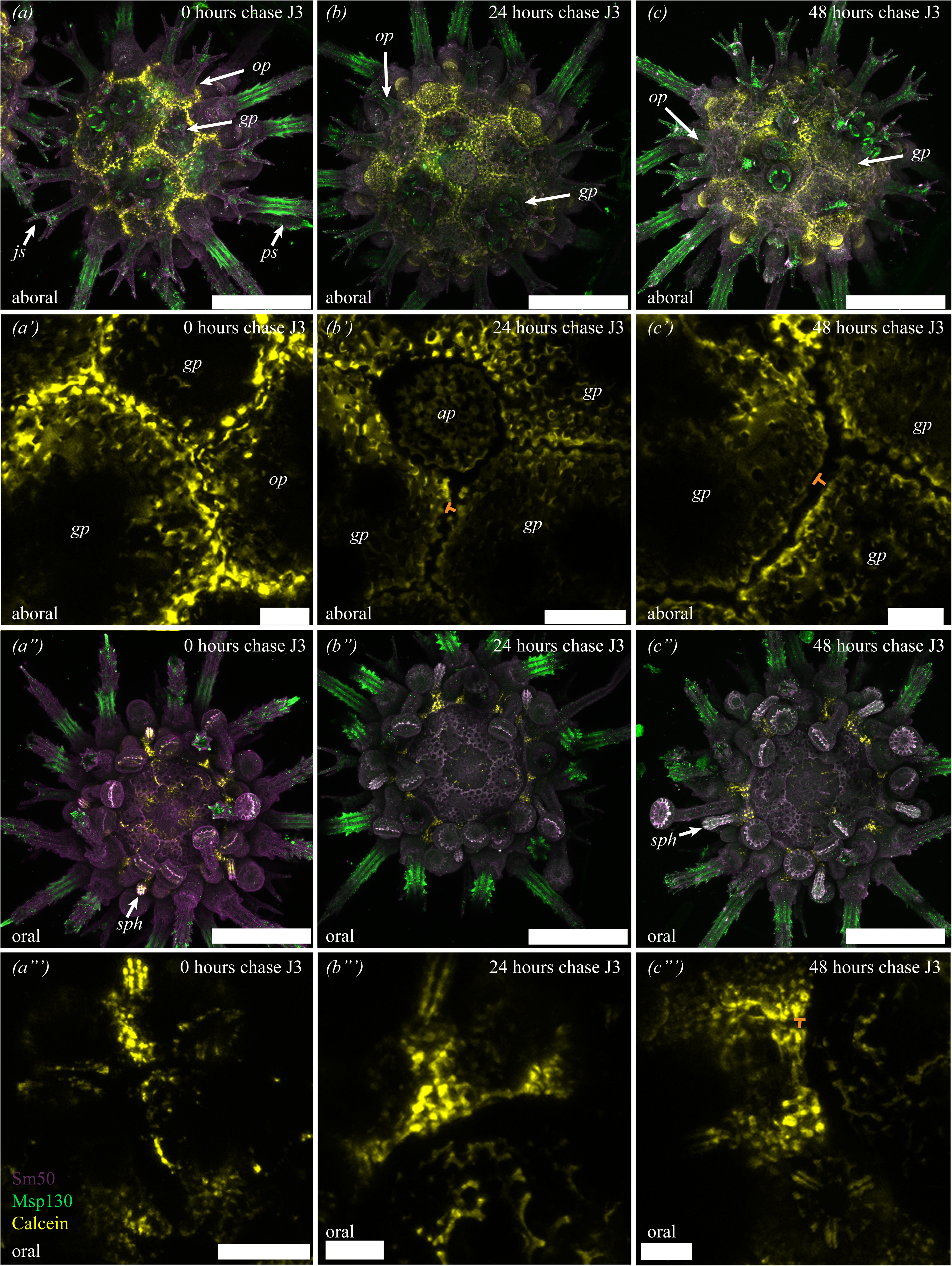
Localization of MSP130 (green), SM50 (purple), and incorporation of calcein (yellow) into growing skeleton in J3 individuals at 0 (a-a’’’), 24 (b-b’’’), and 48 (c-c’’’) hours chase. Description of staining given in the main text. a’, b’, c’ and a’’’, b’’’, and c’’’ are zoomed in images of the above showing incorporation of calcein. Orange bars indicate gap between calcein-marked plated due to subsequent accretion. Abbreviations as in Figure 2 and scale bars in a-c and a’’-c’’ 200 μm, a’,b’’’, c’, c’’ 25 μm, b’ and a’’’ 50 μm.

**Figure 5.**
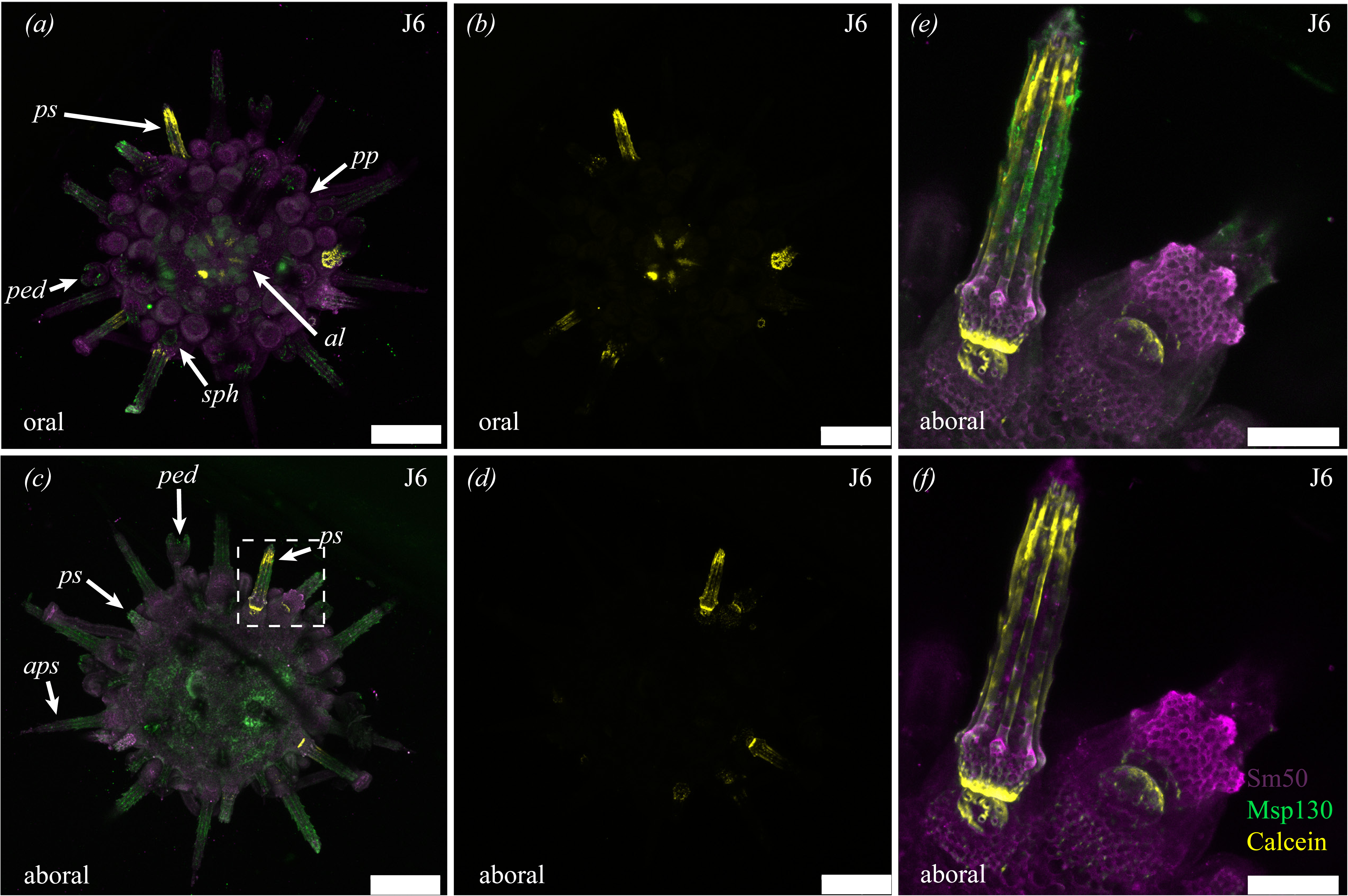
Localization of SM50 (purple), MSP130 (green) and incorporation of calcein (yellow) into J6 animal at 0 hours chase. Description of staining in (a-d) are given in the main text. (e-f) are close ups of the aboral surface showing incorporation of calcein into growing spines and tubercles. Abbreviations as in Figure 2 and scale bars in a-d 200 μm and e-f 50 μm.

**Figure 6.**
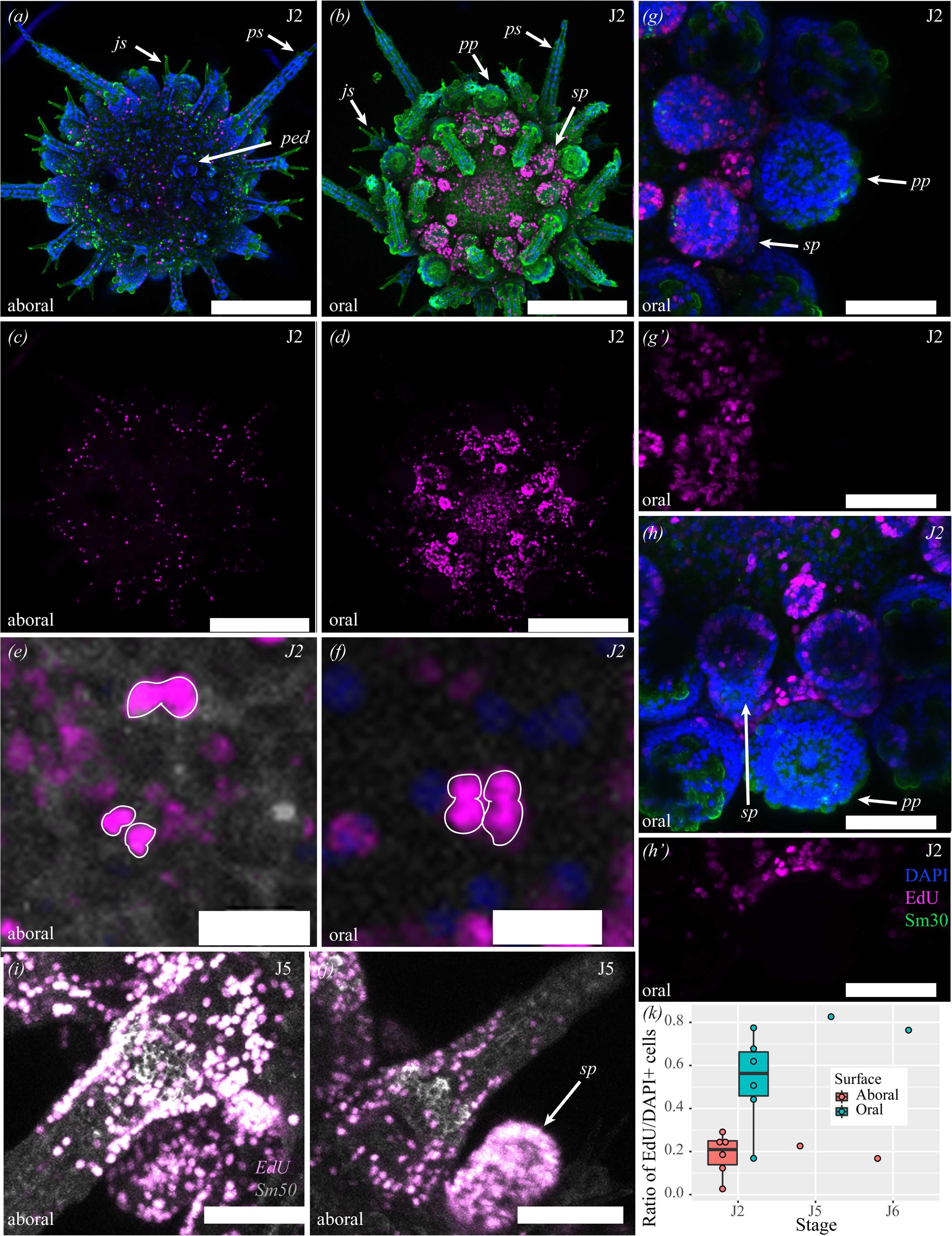
Growth via cellular proliferation in. *P. lividus*. (a-h’) Proliferation on aboral (a,c) and oral (b, d, e, f, g-h’) surfaces in J2 animals. (e-f) close ups of proliferating cell doublets in 24 hour post-chase animal (e) and quadrouplets in 48 hours post-chase animal (f) (g-h’) Zoom showing proliferative zone associated with plate addition. (i-j) Cell proliferation associated with the addition of new ambulacral spines and secondary podia in J5 animal. (k) Graph showing differential cell proliferation on oral and aboral surfaces. Abbreviations as in Figure 2. Scale bars in a-d 200 μm, e-f 10 μm, i-j 100 μm, and g-h 50 μm.

**Figure 7.**
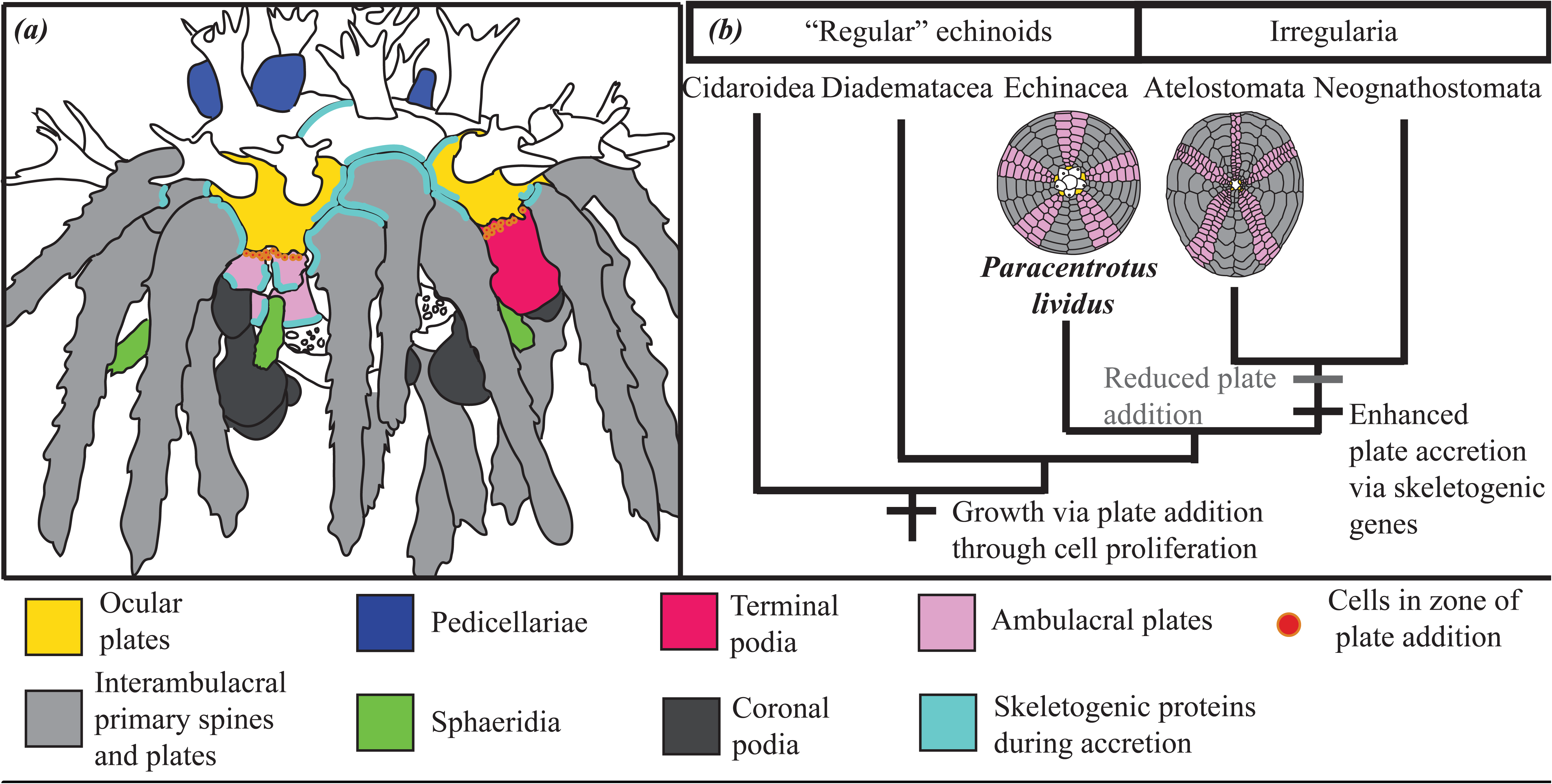
(a) Diagrammatic model showing cell proliferation and expression of skeletal genes during plate accretion and addition in juvenile test growth. (b) Simplified phylogenetic tree of crown group echinoids showing the transitions in sea urchin growth modes in regular and irregular echinoids, and their hypothesized molecular and cellular foundations.

Lastly, to visualize developing skeletal tissues and skeletogenic cells, antibodies were used against three well-characterized proteins known to be involved in echinoid skeletal growth and development. The first of these are the spicule matrix protein 50kDA (SM50) and the spicule matrix protein 30kDA (SM30)[28], two c-lectin type extracellular matrix proteins occluded within the biomineral of the larval and adult skeleton [29-32]. Staining at different stages with both SM30 and SM50 show immunoreactivity with all skeletal structures except for the juvenile jaws and teeth (n= 11;Fig. 2d-g). Crucially, and consistent with differential staining in the larvae and growing juvenile skeleton (Supplemental Fig. 2), staining is stronger in newly formed skeletal elements (identified based on our staging scheme) such as growing ocular plates in J1 individuals and newly added ambulacral spines in J5 animals (Fig. 2f-g). Another marker for skeletal cells is the cell-surface glycoprotein Msp130, known to be specifically expressed skeletogenic cells and skeletal tissues of larval and adult sea urchins [29-31]. The monoclonal antibody 6a9 [33] reacts specifically with Msp130. In different stages in juvenile *P. lividus*, immunostaining with 6a9 identifies all skeletal tissues except for the jaws and teeth (n=10), and in some cases clearly marks cell bodies of skeletogenic cells (as in [25]), which can be seen in the interambulacral spines (Fig.2b, Supplemental Fig. 5). Application of these antibodies in juvenile *P. lividus* has led to a more detailed understanding of the molecular signatures for their musculature, nervous system, and skeleton. In particular, antibodies against skeletal proteins facilitates further study of post-metamorphosis skeletogenesis, and in particular, plate addition and accretion.

### (c) Growth and skeletogenesis of plate accretion

During juvenile growth, skeletal elements like plates and spines, are constantly remodeled and elaborated upon. Plate accretion is the process by which new skeletal tissue is added onto pre-existing skeletal structures [12]. During larval development, cells in the growing portions of the larval skeleton, such as the tips of the arms, express distinct sets of genes (i.e. *Sm30, Sm50* and *Msp130*) relative to other skeletogenic cells [34, 35].We thus hypothesized that plate accretion might similarly involve a distinct set of proteins. To understand the extent to which proteins are differentially or distinctly localized during skeletal growth, and precisely visualize plate accretion, double fluorescent immunostaining of skeletal markers (6a9 specific to MSP130, and anti-SM50) was combined with fluorescent labeling of the growing CaC0_3_ skeleton. In J1 J3 and J6 *P. lividus*, both antibodies are co-localized in all skeletal structures except for the teeth (n=7;Fig. 3a-f). Stronger immunoreactivity for both antibodies was identified in newly formed skeletal tissues, such as the margins of peristomial, ambulacral and interambulacral plates (Fig. 3d-f). The strength of staining of antibodies varies in a tissue-specific manner (Fig. 3d-l), with MSP130 distinctly localized in more distal portions of primary and juvenile spines (Fig. 3j-l) and SM50 more strongly localized in the edges of growing coronal and peristomial plates, in the rosettes of primary and secondary podia, and the bases of newly formed spines in J6 animals (Fig. 3g-l, Fig. 2b). In addition to localizing differentially, SM50 and MSP130 antibodies strongly co-react in the median portions of primary spines (Fig. 2d-f), and co-react less strongly throughout almost all other skeletal tissue. This suggests that these proteins likely co-localize during post-metamorphic skeletal growth.

In addition to immunostaining, calcein [34] was used to identify sites of active skeletogenesis and their distribution relative to the localization of skeletal proteins (Figs. 4, 5). Calcein is a fluorescent marker that binds to Ca^2+^ and is incorporated into the CaCO_3_ skeleton during biomineral deposition [34]. Calcein staining is visible in the sea urchin skeleton following fixation and immunostaining and discrete pulses of incubation with calcein label only skeletal structures growing during the period of incubation. Pulse-chase experiments can thus be used to not only to visualize the position of calcite deposition, but also to determine the spatial extent of subsequent skeletal growth following calcein incubation. J3 (n=5) and J6 (n=1) animals were incubated with calcein for a period of 24 hours, then calcein was washed out and replaced with SW. Animals were fixed and observed immediately after calcein staining (0 hours), or 24 and 48 hours after removal. Figures 4 and 5 show the results of calcein staining in J3 and J6 *P. lividus* juveniles respectively.

After 24 hours of incubation (0 hr chase) in J3 animals, calcein was incorporated into the margins of skeletal structures and plates on both the oral and aboral surfaces. Ocular plates, genital plates, and the anal plate all show calcein incorporated into their growing periphery (Fig. 4a-a’). Calcein was also incorporated into the growing margins of interambulacral, ambulacral, and peristomial plates, as well as in elongating sphaeridia and in the rosettes of growing secondary podia (Fig. 4a’’-a’’’). On the oral surface, growing hemipyramids of the Aristotle’s lantern (the masticatory apparatus) also show strong incorporation of calcein (Fig. 4 a’’’-c’’’, Supplemental Fig. 8-9). In the J6 animal, calcein was incorporated into newly forming ambulacral and interambulacral spines, their corresponding tubercles, as well as in the rosette of a newly added secondary podium (Fig. 5a-b). In J3 animals, the most striking calcein-labeled structures are the growing hemipyramids (Fig. 4a’’), while in J6 animals they are the growing teeth (Fig. 5a-b). This indicates that substantial skeletogenesis is not only taking place in the plates of the test, but also the Aristotle’s lantern prior to the opening of the mouth [25]. Lack of calcein in spines of J3 animals (Fig. 4), but presence in plates, and relative lack of calcein in J6 plates later in development (Fig. 5) suggests modular, piecewise growth. This implied distinct skeletal units grow at distinct times.

In 0 hours chase animals, calcein incorporated into the margins of plates abuts with calcein incorporation in adjacent plates (Fig. 4a-a’). In 24 and 48 hours chase animals, however, there are gaps between the calcein-marked skeleton in adjacent plates, indicating further growth post-incubation (Fig. 4b’, c’). Proximal to distal skeletal growth is also visible in the sphaeridia of 24 and 48 hours chase animals, where calcein is incorporated into the bases of these structure, but not growing tips (Figs. 4b’’, c’’). Newly synthesized skeletal structures marked by calcein, and skeleton deposited post-calcein incubation, show strong immunoreactivity with MSP130 (6a9) and SM50 antibodies (Fig. 4, Supplemental Fig. 9). This can be seen clearly on the aboral surfaces of the animals in Figure 4a-c (Supplemental Fig. 9). This further supports the interpretation that skeletogenic proteins are more abundant in sites of skeletal growth, as is the case in sea urchin larvae [34, 35]. Taken together, these data clarify the morphological changes associated with skeletal growth via accretion and implicate the involvement of skeletogenic matrix proteins. Furthermore, as in the larvae, localization of these proteins suggests that they have both shared and distinct uses in development of the post-metamorphic sea urchin skeleton.

### (d) Cell proliferation in juvenile growth

So far, we have shown that juvenile sea urchin test growth relies on extensive elaboration of pre-existing skeletal structures. Our staging scheme also highlights the addition of new skeletal plates and associated structures such as spines, podia, and pedicellariae during growth. Therefore, it is plausible that cell proliferation might underlie the addition of new morphological structures during the course of echinoid post-metamorphic growth. To identify and quantify proliferating cells during juvenile growth, we used 5-ethynyl-2’-deoxyuridine (EdU), a nucleoside analogue of thymidine incorporated into newly synthesized DNA, to label the nuclei of dividing cells. We then used confocal microscopy to image and quantify EdU^+^ cells on both the oral and aboral surfaces of the test of eights individuals from stages J2, J5 and J6 (Fig. 6). Overwhelmingly, more EdU^+^ cells were present on the oral surface than the aboral surface (Fig. 6a-h, k). To make sure this was not an artifact due to differences in cell densities on each surface, we calculated the ratio of EdU^+^/DAPI-stained nuclei as described in Supplemental Methods (Supplemental Table 2; Supplemental Figure 10). Data clearly show significantly higher ratios of proliferating to non-proliferating cells on the oral surface than the aboral surface (Fig. 6;Mann-Whitney *U* test, *p*=0.0014, *U*=59) a pattern also evident in raw cell counts (Supplemental Table 2).

On the oral surface, EdU^+^ cells are concentrated in growing secondary and peristomial podia (Fig. 6b, d). This contrasts with the primary podia, which form as coelomic outgrowths prior to metamorphosis ([21]; Supplemental Fig 1), and display no evidence of cell proliferation (Fig. 6g-h’). Immediately towards the oral surface of the primary podia and ocular plate, there is a distinct zone of cell proliferation (Fig. 5g-h’). The location of the proliferative zone is crucial, as it corresponds to the position where new plates form during plate addition, and where new secondary podia grow [36]. Pulse-chase experiments (Supplemental Fig. 11) show similar results between 0 and 48 hours chase, though the presence of nuclei doublets and rare quadruplets in 24 and 48 hours chase individuals indicate relatively slow rates of cell division (Fig. 6e-f). Assays of cell proliferation were paired with antibodies against SM30 and SM50, to identify where cell proliferation takes place relative to skeletal growth. There is a general correspondence of EdU^+^ cells to newly grown or growing skeletal structures, such as ambulacral spines in J5 individuals (Fig. 6i-j), Supplemental Fig. 12). There are few EdU^+^ cells at the distal ends of these structures. In J6 individuals, other growing structures, such as new pedicellariae and secondary podia, are associated with an abundance of EdU^+^ cells (Supplemental Fig. 13). This indicates that addition of new morphological structures relies on cell proliferation. The absence of proliferating cells at the distal ends of spines, however, suggests that elaboration and further growth these structures rely less on proliferation, and more on biomineral deposition via differentiated cells, in agreement with our calcein staining (Fig. 5). Assays using EdU thus show that the majority of growth via cell proliferation takes place on the oral surface, while structures on the aboral surface are much less proliferative. Furthermore, immediately below the primary podia there is a high-density of proliferative cells in correspondence with the location of plate addition, suggesting cell proliferation underlies the addition of new test plates of the echinoid test. In addition to new test plates, the growth of new pedicellariae, spines, and tube feet is also associated with high degrees of cell proliferation.

## 4. Discussion

### (a) Adult growth in echinoderms

The growth of juvenile *P. lividus* shows similarities to the growth of other echinoderms, and our experiments shed light on the molecular, cellular, and morphological underpinning of echinoderm growth generally. All eleutherozoans continue to grow throughout their lifetime by adding new axial skeletal elements from a growth zone [10]. It is of interest then, that in just-metamorphosized juveniles of *P. lividus*, the location of the growth zone is located similarly to its position in flattened early post-metamorphic asterozoans. This highlights the fact that the morphology of eleutherozoans begins to diverge extensively after metamorphosis rather than during rudiment development.

Immediately below the ocular plate, our analyses identified a region of proliferating cells coinciding with the zone of plate addition. This zone has been hypothesized as a signalling center responsible for the addition both columns of ambulacral plates, and flanking columns of interambulacral plates, in each ray [10](Fig. 1). Our assays provide insight into the mechanism by which plates are added; namely that the origin of axial tissues formed in each of these paired regions is associated with the proliferation of new cells. Recent knockouts of the pigment genes *Pks* and *Gcm* in the regular echinoid *Hemicentrotus pulcherrimus* support the hypothesis that cell lineages in the echinoid test are partitioned into ten paired regions, each consisting of a half-ambulacrum and half-interambulacrum [37]. Paired with our data on cell proliferation, we hypothesize the existence of 10 discrete aboral proliferative zones, corresponding with each of these paired regions.

### (b) Molecular and cellular underpinning of skeletal growth

Our results suggest that both plate addition and plate accretion contribute to the growth of the sea urchin adult body plan and rely on different molecular and cellular processes (Fig. 7a). Growth at the margins of pre-existing test plates is associated with the intense localization of skeletogenic proteins such as MSP130 and SM50 (Fig. 7a). This is consistent with proteomic analyses that have identified proteins of the MSP130 and SM families in the organic matrix of tests, spines and teeth of adult sea urchins [29, 30]. A previous study using whole mount *in situ* hybridization found the skeletogenic gene *SM37*, the transcription factor *Alx1*, and the signaling molecule *VegfR* expressed in the margins of test plates of juvenile echinoids [38]. In the skeletal cells of embryonic and larval sea urchins, *Alx1* and *VegfR* regulate the expression of skeletogenic genes like *SM30, MSP130* and *SM50* [8]. The precise localization of the skeletogenic proteins MSP130 and SM50 relative to growing skeletal structures in our calcein experiments suggests that upstream regulators such as *Alx1* and *VegfR*, are also important for plate accretion(Fig. 7a).

The addition of new test plates and their associated water vascular and nervous system tissue has been suggested to be analogous to a posterior growth zone as in vertebrates and arthropods [13, 15, 20, 23, 36, 39]. Our results support the idea that the addition of new metameric skeletal structures is associated with cell proliferation. The proliferative zone directly below primary podia is located in the site of plate addition. We can speculate that this proliferative zone may use mechanisms involved in metameric growth across the animal kingdom. Comparative studies have identified Wnt signaling and posterior Hox gene expression as evolutionarily conserved players in posterior growth [40], and such a zone has been hypothesized based on the expression of *Hox11/13* temporally earlier in development of the adult echinoid body plan [13, 15]. Outside of this growth zone, high degrees of cell proliferation in newly added ambulacral spines and pedicellariae indicate that proliferation of cells is also involved in the development of skeletal structures. Indeed, the presence of proliferating cells surrounding structures with strong staining of SM50 suggests that the growth of new anatomical structures may rely on cell proliferation, which may precede skeletal accretion via differentiated cells. Similarly, during regeneration in ophiuroids, differentiated skeletogenic cells which secrete the skeletal plates of the arms are not proliferative [23]. The absence of proliferating cells at the distal tips of newly skeletonizing spines in *P. lividus* suggests a similar scenario.

### (c) Accretion, addition and body plan evolution

Extant echinoids are classified into the globe shaped regular echinoids, which include *P. lividus*, and the irregular echinoids or Irregularia, a clade of bilaterally symmetrical, flattened and oblong forms including sand dollars and sea potatoes (Fig. 7b) [41]. Regular echinoids are paraphyletic with respect to the Irregularia, and the fossil record provides a precise window into the morphological transitions that characterise the evolution of irregular echinoids from their regular echinoid ancestors [42]. While the regular echinoids have displayed marked morphological constraint in their ∼270 million year evolutionary history, the irregular echinoids have drastically diversified their body plans, displaying high phenotypic disparity and heightened rates of morphological evolution [5, 6]. It has been hypothesized that the differential diversity of regular and irregular echinoids is due to differential reliance on the processes of plate addition and accretion during test growth [12]. Regular echinoids rely primarily on continuous new plate addition throughout their lives, while irregular echinoid cease adding plates early in their post-metamorphic ontogeny, and instead grow primarily by accreting onto pre-existing plates [12, 17]. Our experimental results from *P. lividus* allow us to make testable hypotheses concerning the role of cellular processes and gene expression in the evolutionary transition from regular to irregular echinoid body plans. Because irregular echinoids rely less on plate addition, and more on plate accretion, we expect that they grow less via proliferation of new cells. Instead, we hypothesize that they rely more on the activity and expression of skeletogenic genes operating during enhanced plate accretion such as *Sm50* and *Msp130*, to grow their tests (Fig. 7b). In light of our results and given their enhanced reliance on biomineral deposition, we hypothesize that irregular echinoids may have diversified the skeletogenetic toolkit via instances of duplication and/or subfunctionalization of skeletogenic genes throughout their evolutionary history. Future analyses of emerging transcriptomic and genomic datasets will shed light on this hypothesis.

## Supporting information

R code used for cell count quantification

## Acknowledgements

JRT is supported by a Royal Society Newton International Fellowship. PP is supported by Marie Curie ITN EvoCELL (H2020 grant number: 766053 to MIA). Work at the SZN was supported by Assemble+ grant (ASke - 420 – III) to JRT and PO. We would like to thank C. Ettensohn, J. Kohr, R. Burke, and F. Wilt for antibodies and I.F. Berodia, R. Panzuto, and D. Caramiello for assistance culturing sea urchin juveniles and maintenance of adults.

