## Supplementary Materials for "Post-metamorphic skeletal growth in the sea urchin *Paracentrotus lividus* and implications for body plan evolution"

### Supplemental Materials and Methods

#### (a) Culturing

The initial cultures were concentrated at approximately 10 larvae/ml. During the culturing period larvae were then diluted to an approximate final concentration of 1 larva/2ml. They were fed three times a week with 70% *Isocrysis*, 30% *Dunaliela tertiolecta* and aquarium grade fatty acids. The total number of algal cells used varied from 5.000-20,000/ml according to the larval needs as just after microscopic observation of the stomach content. The water changed was twice a week and the larvae were kept at $18^{0}$C. In these conditions the larvae became competent in approximately 3.5 weeks.

#### (b) Immunohistochemistry

Juvenile and late stage *P. lividus* larvae were fixed in 4% paraformaldehyde (PFA) for 15 minutes. Individuals were then treated with ice cold methanol (MeOH) for one minute, and washed four to five times in 1X Phosphate Buffered Saline with 0.1% Tween (PBST). Most washes were for ~5 minutes in 200 μl. Samples were then stored in blocking buffer (BB; PBST with 4% goat or sheep serum and 1% BSA) at 4˚C, either overnight or for up to four months prior to incubation with primary antibodies. Samples were incubated with combinations of primary antibodies for 1.5 hours at 37˚C, or overnight at 4˚C, depending upon the primary antibody. Primary antibody type and dilutions are shown in Supplemental Table 1. Samples were then washed 4 to 5 times with PBST, and incubated with anti-rabbit , anti-rat and/or anti-mouse (Thermo Fischer Scientific) secondary antibodies conjugated with Alexa fluorophores (Supp. Table 1) for one hour using a dilution of 1/1000. Following removal of secondary antibodies, each sample was washed 4 to 5 times with PBST before addition of DAPI at a dilution of 1:10000 (of a stock solution of 5 mg/ml), for at least 15 minutes. To preserve skeleton intact, samples were intentionally not decalcified.

#### (c) Calcein staining

In order to visualize newly deposited $CaC0_3$ of the skeleton, live animals were incubated with Calcein (Sigma) at a dilution of 1:50 of a stock solution of 1.25 mg/ml in sea water. Calcein is light-sensitive, so incubation took place in the dark. Incubation took place in 4 ml wells with a coverslip placed at the bottom of each well. After 24 hours of incubation, calcein was washed out six times with fresh sea water. To understand the spatial incorporation of calcein relative to additional skeletal growth, pulse-chase experiments were carried out following incubation. 0-hours chase animals were fixed immediately following calcein wash out and fixed following protocols for immunostaining up to the BB step and stored at 4˚C in the dark. 24- and 48-hours chase animals were reared in sea water in the dark prior to fixation and storage in BB in the dark at 4˚C. Prior to fixation immunostaining against skeletogenic proteins was carried out as described above, except that all steps were carried out in the dark. Specimens were imaged using an Zeiss LSM 700 confocal microscope.

#### (d) Assays of cell proliferation

To understand the spatial distribution and quantity of proliferating cells during juvenile sea urchin growth assays of cell proliferation were carried out using the Click-iT® EdU Alexa Flour® 555HCS and (Life Technologies) and Click-iT™ EdU Cell Proliferation Kit for Imaging Alexa Flour™ 647 (Thermo Fisher Scientific). J2 animals were incubated with EdU in 7 ml wells at a dilution of 1:1000 (5 μM) in sea water for six hours. J5 and J6 animals were incubated with Edu for 65 hours. Edu was washed out six times following incubation, and 0-hours chase animals were immediately fixed in 4% PFA for 15 minutes, while 24-and 48 hours chase animals were further reared in wells until fixation. After fixation, samples were washed twice with PBST, and permeabilized in PBST+Triton for 30 minutes. They were then washed twice in PBST, permeabilized in MEOH for 1 minute, and washed a further two times before storage in BB overnight at 4˚C. Primary and secondary antibody incubations were carried out as described above, and following the PBST washes after secondary antibody incubation, samples were incubated in 50 μl of the Click-iT® Reaction cocktail, following the manufacturer's instructions for 30 minutes. Samples were then washed with 50μl of Click-iT® reaction Rinse Buffer for 30 minutes, before two washes in PBST, and addition of DAPI at a dilution of 1:10000 (of stock a solution of 5 mg/ml). Specimens were imaged using an Zeiss LSM 700 confocal microscope. For each image, five circles of identical size were used to define regions of interest (ROIs) on each image. The relative position of the circles on each image was not standard with respect to the image, thus resulting in a semi-random placement of each circle. This was done intentionally so as to not bias which portion of the animal surfaces were counted. $EdU^+$ and $DAPI^+$ nuclei were quantified in ImageJ by first converting images to 16-bit, then thresholding. Images were then converted to binary images using Process→Binary, and then Process→ Binary →Watershed. Finally cells within the ROIs were counted using Analyze→Analyze Particles. This is summarized in Supplemental Figure 11. The ratio of $Edu^+$ to $DAPI^-$stained nuclei was then counted for each image, and the ratios for the aboral and oral surfaces of each quantified individual were compared using a Mann-Whitney *U* test. R code to repeat analyses and plotting are in Supplemental File 1.

**Supplemental Data**

**(1) Staging Scheme**

In order to make standardized comparisons across cultures and asynchronously developing individuals, we devised a staging scheme based off the presence and absence of specific morphological characters during the course of post-metamorphic development. Our staging scheme does not cover the most immediate stages of post-metamorphic growth, and for an account of those stages we refer the reader to Gosselin and Jangoux [1]

(i) Stage Juvenile 1 (J1) —The earliest of our stages, J1 corresponds to approximately 1-3 days post-metamorphosis. At this stage, the post-metamorphic juvenile is readily characterized by the primary podia, and the interambulacral primary spines. There are four interambulacral spines in each interambulacral area, one on each interambulacral plate. Ambulacral regions are dominated by the primary podia, and two poorly developed secondary podia are present in each ambulacral area. The primary podia have well-formed disks and rosettes, while the secondary podia lack disks, and have not yet formed rosettes. There are two small ambulacral plates in each ambulacrum, and there are ten peristomial plates arranged inter-radially adoral to each interambulacrum. When viewed adorally, the five ocular plates, five genital plates, and an anal plate are visible, and two juveniles spines (or multified spines of Gosselin and Jangoux [1]) are present on each ocular plate. The ocular plates each bear a ledge, which extends slightly over top of the primary podia. Between one and two pedicellariae are present on two of the genital plates, and at least one juvenile spine is present on each genital plate. The plates of the periproct and genital plates do not appear well-sutured together.

(ii) Stage Juvenile 2 (J2) —J2 animals correspond to approximately 4-7 days post-metamorphosis. This stage shows the additional growth on the oral surface relative to J1 individuals, and is primarily differentiated based upon the presence of five peristomial tube feet, and the development of disks and rosettes in the secondary podia of each ambulacrum. There are still four primary spines and interambulacral plates present in each interambulacrum. Ambulacral areas are still dominated by the primary podia, though secondary podia are much larger, about the same size as the primary podia. Secondary podia have well-developed disks and growing skeletal rosettes. Small, peristomial podia are present protruding through five of the peristomial plates. Ocular plates still bear two juvenile spines, and there are between one and two pedicellariae on two genital plates. Genital plates do not appear well-sutured.

(iii) Stage Juvenile 3 (J3) —J3 stage animals are roughly 7-10 days post-metamorphosis. They are most readily characterized by the presence of newly added pedicellariae, spine-like sensory structures present along one of the most oral ambulacral plates in each ambulacrum. All plates of the test have grown larger at their margins, and interambulacral areas bear four plates and primary spines. Ambulacral plates have grown to abut one another and the interambulacral plates and a clear circular margin has formed around the peristome. The secondary podia bear well-developed disks and rosettes, and are about the same size as, and may appear larger than, the primary podia. Spine-like, elongate sphaeridia have started to grow towards the perradial margin of one ambulacral plate in each ambulacrum, but they lack the large bulb present in later stages. Peristomial podia have developed small disks. On the aboral surface, the genital and ocular plates have started to fuse via plate accretion, and the anal plate has grown larger and more distinct. The number of juvenile spines and pedicellariae remains unchanged from earlier stages.

(iv) Stage Juvenile 4 (J4) — Stage J4 animals are approximately 8-12 days post-metamorphosis. They are differentiated from Stage J3 animals by having a well-developed bulb at the tip of sphaeridia, and by having begun to add a second row of secondary podia. Interambulacra consist of four plates, each with a primary spine. Ambulacra are dominated by secondary podia and sphaeridia. Sphaeridia are elongate and with a well-developed bulb at their tip. Aboral surface shows well-sutured genital and ocular plates.

(v) Stage Juvenile 5 (J5) —Stage J5 animals are substantially older than earlier stages, being approximately 2.5-3.5 weeks post-metamorphosis. This gap in timing is reflected by the addition of numerous novel morphological structures relative to earlier stage animals. In J5 animals, the aboral surface is relatively smaller than the oral surface, and all news growth has taken place in axial tissues. There are still four interambulacral plates and primary spines in each interambulacral area, but interambulacral pedicellariae have been added to the most aboral interambulacral plate in each area. Ambulacral areas have well-developed, bulbous and glossy pedicellariae, and primary podia are no longer visible, presumably having been lost during the course of growth. There are two well-developed rows of secondary podia in each ambulacral area, with each podium bearing a disk and rosette. One or two secondary podia of the third row are growing adjacent to the ocular plate, but have yet to develop disks. Immediately aboral of the ocular plate, ambulacral areas are dominated by a large ambulacral primary spine. In the extraxial tissues on the aboral surface, very little growth has taken place. Juvenile spines are still present on the aboral surface, but they are not easily visible. By J5, juveniles have also opened their mouths.

(vi) Stage Juvenile 6 (J6) —The final stage in our scheme, J6, corresponds to roughly 3-4 weeks post-metamorphosis. J6 animals are the most morphologically complex stage, being characterized primarily by the addition of interambulacral and ambulacral primary spines, as well as the addition of ambulacral pedicellariae adoral to the ambulacral primary spines. There are four or five interambulacral plates in J6 animals, however, the plate surfaces have grown, and new interambulacral primary spines have been added to the adapical most interambulacral plates in each area. There are thus five or six interambulacral primary spines in each interambulacrum. As in J5 animals, a single pedicellariae is also present in each interambulacral area. Pedicellariae have also formed on a single ambulacral plates adoral to the largest ambulacral primary spines (which was added in J5 animals) in some of the ambulacra. Well-developed sphaeridia are present in each ambulacrum. Like J5 animals, J6 animals have no primary podia, and their adoral ambualcra are dominated by ambulacral primary spines. There are two ambulacral primary spines in each ambulacrum. There are three well-developed rows of secondary podia in each ambulacrum, and the podia of a fourth row are growing. Like J5 animals, no growth of new structures is taking place in the extraxial tissues of the abora surface.

**Supplemental Figure Captions**

Fig. S1 Confocal images of pre-metamorphosis, late stage *P. lividus* larvae showing developing adult body plan within the rudiment stained for MSP130 using monoclonal antibody 6a9 (green), and SM50 using an anti-SM50 antibody (magenta). (a) Merged image of Sm50 and Msp130 and nuclear staining (DAPI, blue) showing developing skeletal tissues, within rudiment including spines, tubercles, and the rosette of a primary podium. DAPI (blue) Msp130 strongly labels the bodies and projections of cells tightly associated with the skeleton (skeletogenic cells), and the skeleton. Conversely, staining for SM50 is stronger in the skeleton itself, and in structures such as the spines and rosette. (b) Interpretive drawing showing morphology corresponding to (a). (c) Magenta channel showing staining for SM50. Stronger staining of SM50 is identified at the base of te spines and rosette, where deposition of skeletal tissue is active. In contrast, the larval skeleton is not strongly labeled. (d) Green channel showing staining for MSP130 using monoclonal antibody 6a9. Note the strong staining in the larval skeleton, and the staining of the skeletogenic cells. sc, skeletogenic cells; ls, Larval skeleton; pp, primary podium; ros, rosette. Scale bar is 50 μm.

Fig. S2. Post-metamorphic *P. lividus* juveniles. Light microscope images showing post-metamorphis juvenile *P. lividus*. (a) Oral view of stage J4 juvenile. Sphaeridium is well developed and oral surface is dominated by tube feet. Pigmented cells are abundant in the distal disk of the tube feet. (b) Lateral view of J5 animal. Animal is oblong in lateral view, and the aboral surface bears far fewer structures than the oral surface. (c) Oral view of same animal as (b). Note well-developed, circular peristomial margin. Pedicellariae are present on interambulacral plates, and well-developed, glossy sphaeridia are present on ambulacral plates in addition to ambulacral spines. (d) Aboral surface as same animal in (b)-(c). The aboral surface bears few structures, except for the juvenile spines present on genital and ocular plates. Darkened crescent is the gut, visible inside of the animal. sp, secondary podia; sph, sphaeridia; ps, primary interambulacral spine; js, juvenile spine; ped, pedicellariae; as, ambulacral spine.

Fig. S3. Skeleton and Musculature of *P. lividus* juveniles. Light sheet microscope images showing staining in J1 individual for Myosin Heavy Chain (MHC) using an anti-MHC antibody and MSP130 using monoclonal antibody 6a9. (a) Lateral view showing staining for MHC (Magenta), MSP130 (Green), and nuclei using DAPI (Blue). MSP130 strongly stains the interambulacral primary spines and the juvenile spines, while MHC is localized in the muscular tissue of the primary podia, in longitudinal bands surrounding the tubercles, in bundles within the spines, and in the bases of the pedicellariae pincers. (a') Magenta channel from (a) showing only MHC. (a'') Magenta and blue channels showing MHC and DAPI. Staining for MHC is strongest in the bases of the pedicellariae. (b) Close up of primary interambulacral spine showing bundles of $MHC^+$ cells within the lumen of skeletal spines. (b') Magenta channel from (b), showing bundles of $MHC^+$ cells. (b'') Bundles of $MHC^+$ cells relative to DAPI staining, indicating that each $MHC^+$ circle stains multiple cells as opposed to a single cell. (c) Enlargement of pedicellarial staining. The most intense staining for MHC in the animal is the base of the pedicellariae pincers. (d) Close up from (a) with only magenta and green channels showing longitudinal bands of $MHC^+$ cells surrounding the tubercles of primary interambulacral spines and muscles along the primary podia. ps, primary interambulacral spine; js, juvenile spine; pp, primary podia; ped, pedicellariae. Scale bars in (a)-(a'') are 100 μm, rest are 25 μm.

Fig. S4 Localization of neuronal markers in early juveniles. Confocal microscope images showing staining in J1 individual for synaptotagmin using 1E11 and serotonin using an anti-serotonin antibody. (a) Oral view of J1 individual showing immunoreactivity for synaptotagmin (green), and serotonin (magenta). Nuclei are stained with DAPI (blue). Synaptotagmin is localized within the interior of the primary podia, and in bands surrounding the tubercles and bases of the spines. Synaptotagmin and serotonin co-localize in single cells arranged proximo-distally along the lumen of the primary interambulacral spines (b) Close up of (a). (c) Aboral view of same individual as (a). (d) Green channel from (a) showing synaptotagmin. (e) Green channel from (b) showing synaptotagmin. Note the $synaptotagmin^+$ cells lining the lumen of the primary interambulacral spines. (f) Green channel from c showing synaptotagmin. (g) Magenta channel from a showing serotonin. (h) Magenta channel from (b) showing serotonin. Note the $synaptotagmin^+$ cells lining the lumen of the primary interambulacral spines, which we interpret to be serotonergic neurons. (i) Magenta channel from (c) showing Serotonin. (j) Green and blue channels from (a) showing synaptotagmin and DAPI. (k) Green and blue channels from (b) showing synaptotagmin and DAPI. (l) Green and blue channels from (c) showing synaptotagmin and DAPI. All scale bars in (c), (f), (i), (l) are 50 μm, rest are 100 μm.

Fig. S5. Neuronal and skeletal markers in early juveniles. Confocal images showing localization of ELAV and MSP130 in J2 individual. (a) Staining for ELAV using an anti-ELAV antibody

(Magenta), MSP130 using monoclonal antibody 6a9 (Green), and DAPI (blue). MSP130 is strongly localized in the primary and juvenile spines, and the rosettes while ELAV is localized in cells in the primary and secondary podia. (b) Close up of same invididual as (a) showing ELAV$^+$ cells, interpreted to be neurons, in the secondary podia. In the interambulacral primary spines, MSP130 is localized in the cell bodies of cells that we interpret to be skeletogenic cells. (c) Magenta and green channels of (a), showing staining of ELAV and MSP130. (d) Magenta and green channels of (b). Importantly MSP130 and ELAV are differentially localized in the secondary podia, with ELAV marking neurons, and MSP130 staining the growing rosette. (e) Magenta and blue channels of (a), showing ELAV and DAPI. (f) Magenta and blue channels of (b), showing ELAV and DAPI. Cell nuclei are marked by DAPI, but ELAV stains the surrounding cell body. (g) Green and blue channels of a, showing MSP130 and DAPI. (h) Green and blue channels of (b), showing MSP130 and DAPI. As with ELAV, MSP130 stains the cell body, while nuclei of the same cells are marked with DAPI. n, neuron; ros, rosette; sc, skeletogenic cell. Scale bars in (a), (c), (e), (g) are 200 μm, scale bars in (b), (d), (f), (h) are 100 μm.

Fig. S6. Specific and pan-neuronal markers. Confocal images showing localization of ELAV and Synaptotagmin in J1 individual. (a) Staining for synaptotagmin (green). Notice rings of nerves surrounding tubercles and in bundles of neurons in the primary spines. (b) Staining for ELAV. Notice bundles of neurons in primary spines. (c) DAPI staining showing distribution of cell nuclei. (d) Merge showing co-localization of Synaptotagmin and ELAV. Of importance is the fact that co-localization is most strong in the bundles of neurons at the base of the primary spines. Scale bars in (a)-(c) are 200 μm, (d) is 50 μm.

Fig. S7 Localization of acetylated tubulin. Confocal images showing staining for acetylated tubulin and DAPI in J1 animals. (a) Merge showing staining for acetylated tubulin (green) using an anti-acetylated tubulin antibody and for cell nuclei using DAPI (blue) on the oral surface. Cilia cover the entire surface of the animal and the lateral edges of the spines, as shown in Gosselin and Jangoux [1]. Acetylated tubulin is also found in the interior of the primary and secondary podia, which we interpret to be tissue of the nervous system in agreement with a recent study [2]. (b) Staining for acetylated tubulin and cell nuclei on the aboral surface of a different J1 individual. Cilia are found across the surface of the animal, and show a high density on pedicellariae. The individual from (a) is shown in the bottom of the image. (c) Green channel from a showing only staining for acetylated tubulin. Note the cilia lining the margin of the discs of the primary podia. (d) Green channel from (b) showing staining of acetylated tubulin. pp, primary podia; sp, secondary podia, ps, primary interambulacral spine; ped, pedicellariae. Scale bar in all images is 200 μm.

Fig. S8 Calcein localization relative to skeletal proteins. Confocal microscopy showing staining of SM50 using an anti-SM50 antibody (magenta), and MSP130 using monoclonal antibody 6a9 (green), as well as the incorporation of fluorescent calcein (yellow) into the growing J3 sea urchin test. (a) Aboral surface showing incorporation of calcein into the margins of accreting genital, anal, ocular and interambulacral plates. MSP130 and SM50 are also shown, and reveal distinct patterns of localization and co-localization. (b) Oral surface of same individual shown in (a). Calcein is shown incorporated into margins of accreting ambulacral and interambulacral plates, elongating sphaeridia, growing hemipyramids, and rosettes. There is co-localization of

SM50 and msp130 in sites of calcein incorporation, implicating SM50 and MSP130 in active skeletogenesis. (c) Same as (a), with calcein removed. Co-localizaion of MSP130 and SM50 is indicated by greyish-white color. This is evident at the margins of genital and interambulacral plates, as well as in the tubercles. Compared with (a) and (e), it is evident that strong co-localization of these skeletogenic genes occurs in sites of accretion as identified using calcein. (d) Same as (b), with calcein removed. Co-localization of MSP130 and SM50 is evident in the margins of accreting peristomial, ambulacral, and interambulacral plates, as well as in the elongating sphaeridia and rosettes of secondary podia. Compared with (a), and (f), this co-localization is evident in sites of calcein incorporation. (e) Yellow channel from (a), showing sites of calcein incorporation in genital, interambulacral and anal plate margins. Compare with co-localization of skeletogenic proteins in (c). (f) Yellow channel from (b), showing sites of calcein incorporation in margins of accreting peristomial, ambulacral, and interambulacral plates, and growing hemipyramids, rosettes and sphaeridia. Compare with sites of skeletogenic protein localization in (d), (h), and (j). (g) Purple channel from (a), showing localization of SM50 protein. Immunoreactivity is stronger in the bases of spines and tubercles, and in the margins of genital, anal and interambulacral plates. (h) Purple channel from (b), showing localization of SM50. Immunoreactivity is stronger in margins of peristomial plates, elongating sphaeridia, and rosettes of secondary podia. (i) Green channel from a showing localization of MSP130 proteins using 6a9. Immunoreactivity is stronger in the medial and distal portions of interambulacral spines, and in the distal portions of juvenile spines and pedicellariae. (j) Green channel from (b) showing localization of MSP130 protein using 6a9. Localization is strongest in medial and distal portions of interambulacral spines, in the rosettes, in elongating sphaeridia and in the margins of peristomial plates. ps primary spine; js, juvenile spine; ped, pedicellariae; sph, sphaeridia, pp, primary podia; sp, secondary podia; hp, hemipyramid. Scale bar in all images is 200 μm.

Fig. S9. Skeletal growth over time. Confocal microscopy showing staining for SM50 using an anti-SM50 antibody (magenta), and the incorporation of fluorescent calcein (yellow) into the J3 sea urchin test at 0, 24, and 48 hours chase. Specimens are the same as in Figure 4 in the main text, with green channel (MSP130) removed. (a) Aboral surface from 0 hours chase animal showing incorporation of calcein) relative to localization of SM50. SM50 is localized throughout all visible skeletal structures, including localization in areas of calcein incorporation such as the margins of plates, shown in whiteish-grey. (a’) Yellow channel from (a), showing sites of calcein incorporation in aboral surface of 0 hours chase animal. (a’’) Oral surface from 0 hours chase animal, showing incorporation of calcein into skeleton relative to localization of SM50. SM50 stains the entire skeleton except for the teeth, and is localized in sites of calcein incorporation in the sphaeridia, rosettes, and margins of peristomial, ambulacral, and interambulacral plates. (a’’’) Yellow channel from (a’’) showing incorporation into oral surface of 0 hours chase animal. (b) Aboral surface of 24 hours chase animal showing incorporation of calcein relative to staining for SM50. Additional skeletogenesis at the margin of plates has taken place in the 24 hours since calcein incubation, as evident in the gaps between calcein-incorporated skeleton in adjacent plates. SM50 is still strongly localized in some areas of calcein incorporation, such as the tubercles and margins of plates. (b’) Yellow channel from (b), showing incorporation of calcein into aboral surface of 24 hours chase animal. (b’’) Incorporation of calcein and localization of SM50 protein in oral surface of 24 hours chase animal. SM50 is localized where calcein was incorporated into the skeleton in the ambulacral, peristomial and interambulacral plates, but strong SM50 localization is also evident in skeleton that has been deposited subsequent to incubation with calcein. This is most evident in the margins of the peristomial plates, in the elongating sphaeridia, and in the rosettes of secondary podia. (b''') Yellow channel showing incorporation of calcein in the skeleton of the oral surface of a 24 hours chase animal. (c) Incorporation of calcein and localization of SM50 in the skeleton of the aboral surface of a 48 hours chase animal. Gaps between sites of calcein incorporation in adjacent plates (of maximum size of 7.5 μm) are clearly visible on the aboral surface. Additionally, strong SM50 localization in these gaps, and in other structures such as the tubercles, bases of primary spines, and margins of plates are indicative of further biomineralization involving SM50. (c') Yellow channel from c showing incorporation of calcain into aboral skeleton of 48 hours chase animal. (c'') Oral surface of 48 hours chase animal showing incorporation of calcein and localization of SM50. Substantial growth has taken place in the 48 hours since the incorporation of calcein, which is evident in the strong localization of SM50 in the sphaeridia, in the margins of the peristomial plates, and in the rosettes of secondary podia. Additionally, gaps between areas of calcein incorporation are present in ambulacral and interambulacral plates. (c''') Yellow channel of (c'') showing incorporation of calcein into oral surface of 48 hours chase animal. iamb, interambulacral plate; op, ocular plate; sph, sphaeridia; pp, primary podia; sp, secondary podia. Scale bar in all images is 200 μm.

Fig S10. Images showing different stages of the strategy used to quantify $EdU^+$ nuclei relative to all nuclei. (a) and (b) Show raw images for $Edu^+$ (a) and $DAPI^+$ cells (b). (c-d) Binary images for quantification. (e-f) Five regions of interest s of binary cells for quantification. See expanded details in Supplemental Methods.

Fig. S11. Cell proliferation in early juvenile stages. Confocal microscope images showing the incorporation of EdU into the oral surface of growing J2 sea urchins, as well as staining for SM30 using an anti-SM30 antibody. (a) Staining for EdU (magenta), SM30 (gray), and cell nuclei using DAPI (blue) in the oral surface of a 0 hours chase individual. Most proliferation takes place in the secondary podia and growing peristomial podia. Additionally, immediately adoral to the primary podia, there is a zone of cell proliferation corresponding to the zone where new plates are added. Of interest, there is no proliferation in the primary podia themselves, which form in the rudiment and atrophy within a few weeks following metamorphosis [1]. (a') Gray channel from a showing localization of SM30. Protein is localized in all skeletal tissue except for the teeth and is most strongly localized in the primary interambulacral spines. (a'') Localization of SM30 relative to proliferating cells marked by EdU. There is very little proliferation in the primary spines and in the peristomial test plates. (b) Staining for EdU, SM30 , and cell nuclei using DAPI in a 24 hours chase individual. As in the 0 hours chase individuals, most proliferating cells are located in the secondary and peristomial podia, and the proliferative zone adoral to the primary podia. (b') Gray channel from (b), showing localization of SM30. (b'') Localization of SM30 relative to proliferative cells marked by EdU. (c) Staining for EdU, SM30, and cell nuclei using DAPI in a 48 hours chase individual. Localization of $EdU^+$ cells does not differ substantially from 0 and 24 hours chase individuals. Presence of cell doublets indicates that rates of cell division are relatively slow, and that not much cell division has taken place in the 48 hours since incubation with EdU. (c') Gray channel showing localization of SM30. (c'') Location of $EdU^+$ cells relative to skeletal tissues identified using SM30. ps, primary interambulacral spine; pp, primary podia; sp, secondary podia. Scale bar in all images is 200 μm.

Fig. S12. Cell proliferation in J5 Juvenile. Incorporation of EdU into a 0 hours chase J5 individual after 63 hours of incubation with EdU, and staining for SM30 using an anti-SM30 antibody. (a) Aboral surface of 0 hours chase animal showing localization of SM50 relative to proliferating cells marked by EdU (magenta-white). There is relatively little cell proliferation present in extraxial tissues on the aboral surface such as the genital and periproctal (anal) plates. Most cell proliferation seen from this view is associated with novel structures, such as newly added secondary podia and ambulacral spines. (b) Proliferating cells marked by EdU and localization of SM50. Proliferation is extensive on the oral surface in axial tissues, especially when compared to the extraxial tissues of the aboral surface. High degrees of cell proliferation are found in the disks of the secondary podia, as well as in the peristomial podia and sphaeridia. Most proliferating cells on spines are located more proximally towards the base of the spine. (c) Magenta channel from (a), showing location of proliferating cells on aboral surface. (d) Magenta channel from (b), showing location of proliferating cells on oral surface. (e) Close up view of cell proliferation associated with growth of a newly added ambulacral primary spine. Most proliferating cells are localized towards the base of the spine and the tubercle. Strong staining of SM50 is also located towards the base of the spine, though we interpret most distal growth takes place via skeletogenesis. (f) Cell proliferation associated with a growing primary spine and secondary podia. As in (e), most proliferating cells are located nearer to the base of the spine. High degrees of cell proliferation are associated with the growth of the secondary podia, as is also seen in J2 individuals in Figure S11. (g) On the oral surface, high degrees of cell proliferation are associated with the tube feet. In particular, the margins of the tube feet disk show high degrees of cell proliferation, potentially associated with sensory motor neurons. ps, primary interambulacral spine; aps, ambulacral primary spine; sp, secondary podia; ped, pedicellariae; m, mouth. Scale bars in (a)-(d) are 200 μm, scale bars in (e)-(g) are 100 μm.

Fig. S13. Cell proliferation in a J6 Juvenile. Incorporation of EdU into a 48 hours chase growing J6 sea urchin after 63 hours of incubation with EdU. (a) EdU stained nuclei (magenta) relative to all cell nuclei marked with DAPI (gray) on the aboral surface. Note the low abundance of proliferating cells on extraxial tissue (genital and anal plates). Most cell proliferation in this view is associated with the addition of new structures, such as ambulacral and interambulacral primary spines. (b) Cell proliferation marked using EdU on the oral surface of a J6 animal. High degrees of cell proliferation are associated with newly added pedicellariae. Proliferating cells are also found in the circular margins of the disks of the secondary podia. (c) Magenta channel of (a), showing location of cells marked with EdU. (d) Magenta channel of (b), showing EdU marked cells on oral surface. (e) Close up of newly added ambulacral and interambulacral spines, secondary podia, and associated cell proliferation. (f) Close up of proliferating cells in newly added secondary podia and spines. (g) Close up of ambulacral pedicellariae and associated cell proliferation. ps, primary interambulacral spine; aps, ambulacral primary spine; sp, secondary podia; ped, pedicellariae; m, mouth. Scale bars in (a)-(d) are 200 μm, scale bars in (e)-(g) are 100 μm.

**Supplemental Table Descriptions**

Table S1. Antibodies and dilutions used in this study.

Table S2. Quantification of EdU+ nuclei relative to all nuclei labeled with DAPI

[2] Formery, L., Orange, F., Formery, A., Yaguchi, S., Lowe, C.J., Schubert, M. & Croce, J.C. Neural anatomy of echinoid early juveniles and comparison of nervous system organization in echinoderms. *Journal of Comparative Neurology*.

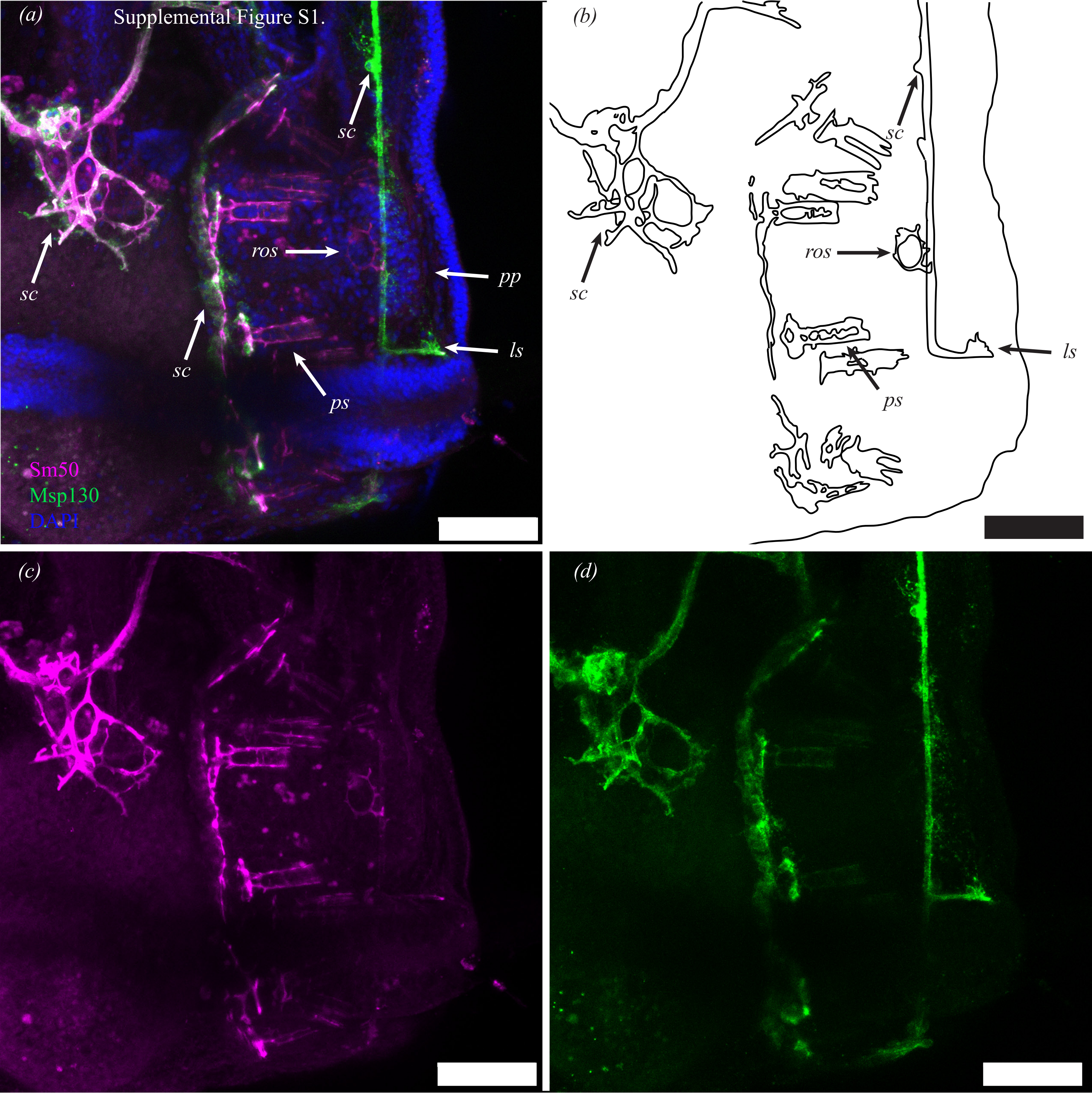
Supplemental Figure S1.
(a)
sc
sc
sc
ros
pp
ls
ps
Sm50
Msp130
DAPI
(b)
sc
sc
ros
ls
ps
(c)
(d)

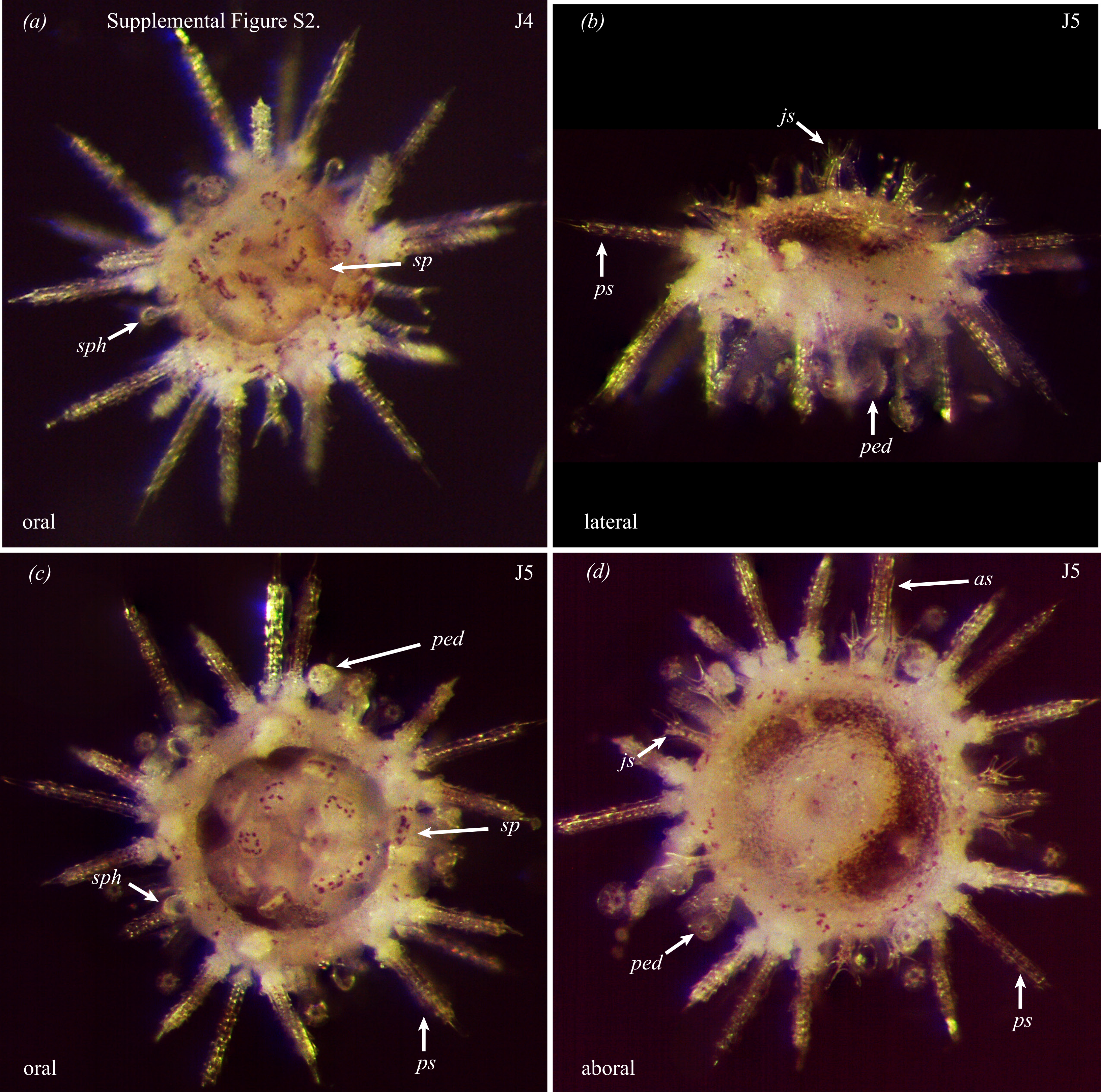

(a)
Supplemental Figure S2.
J4
sp
sph
oral
(b)
J5
js
ps
ped
lateral
(c)
J5
ped
sp
sph
ps
oral
(d)
J5
as
js
ped
ps
aboral

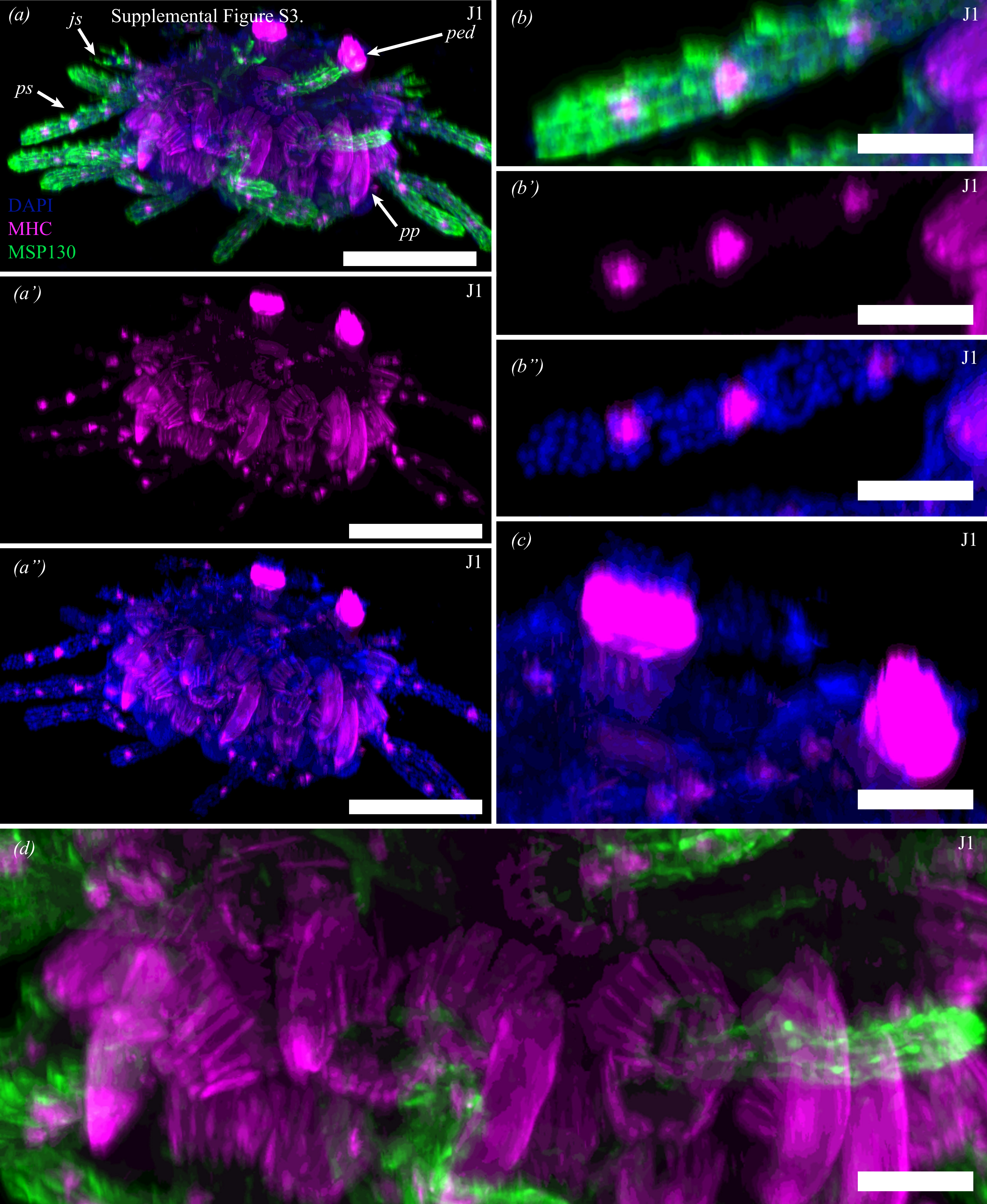
Supplemental Figure S3.
(a)
js
ped
ps
pp
DAPI
MHC
MSP130
J1
(a')
J1
(a'')
J1
(b)
J1
(b')
J1
(b'')
J1
(c)
J1
(d)
J1

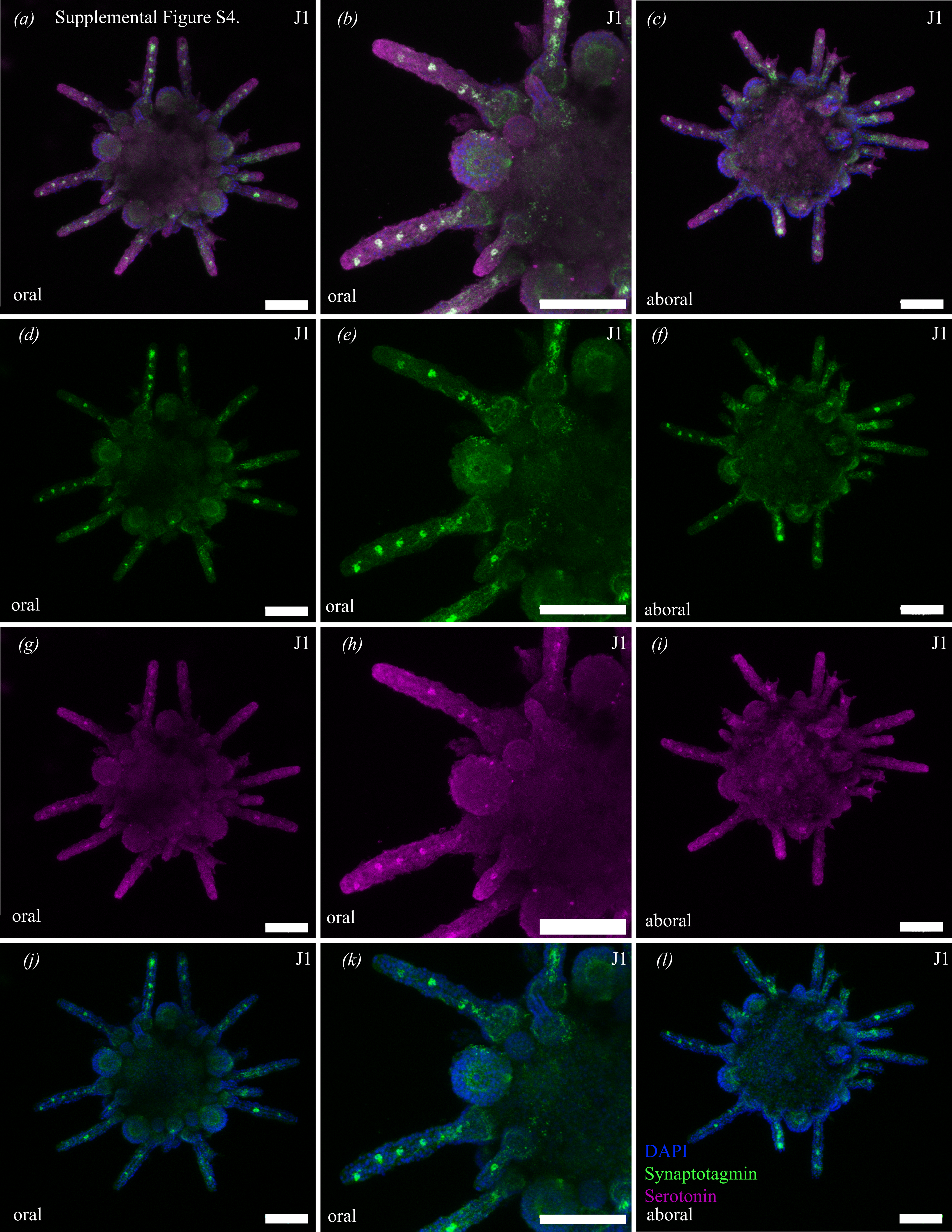

(a)
Supplemental Figure S4.
J1
oral
(b)
J1
oral
(c)
J1
aboral
(d)
J1
oral
(e)
J1
oral
(f)
J1
aboral
(g)
J1
oral
(h)
J1
oral
(i)
J1
aboral
(j)
J1
oral
(k)
J1
oral
(l)
J1
DAPI
Synaptotagmin
Serotonin
aboral

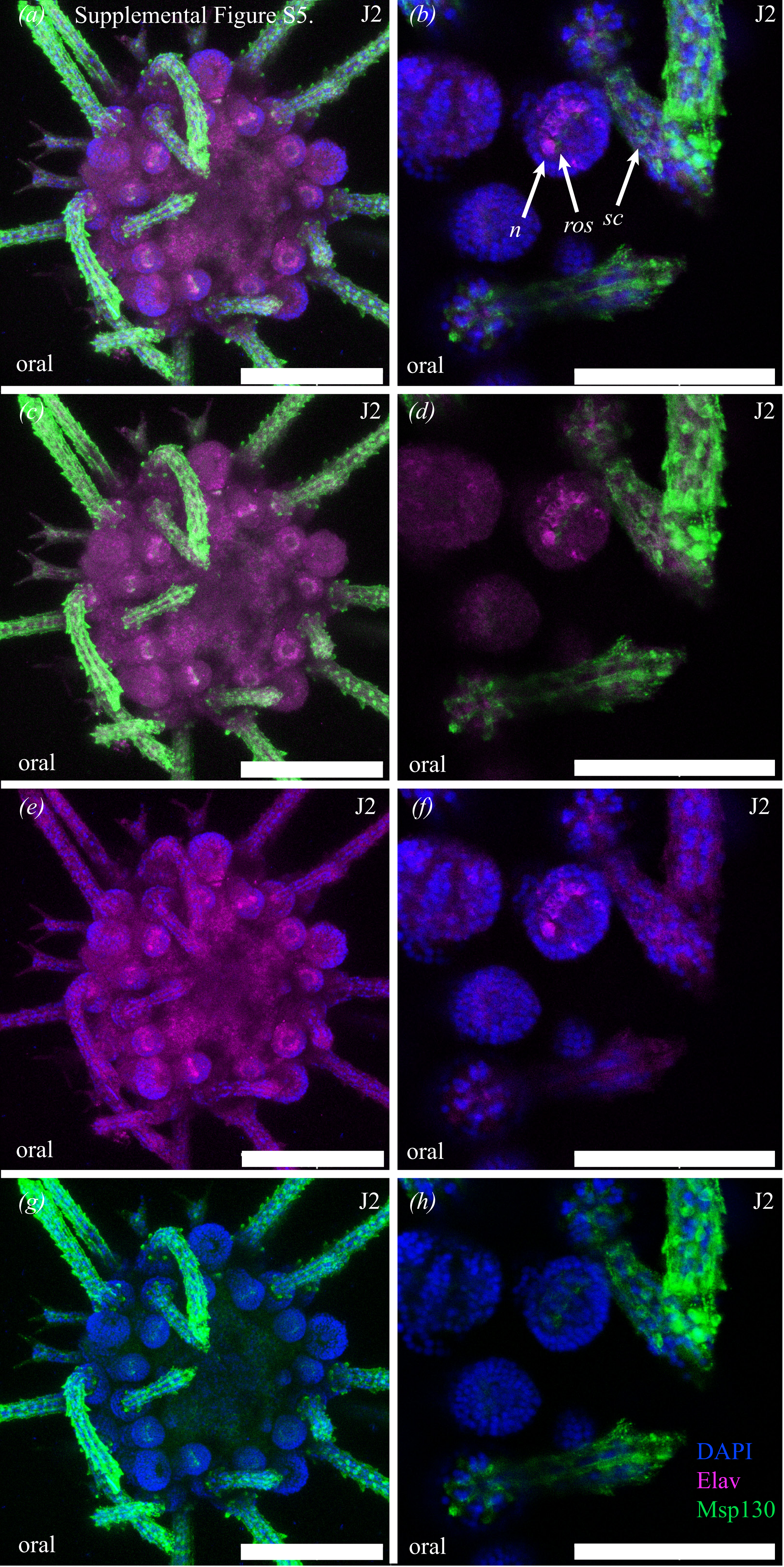
(a)
Supplemental Figure S5.
J2
oral
(b)
J2
n
ros
sc
oral
(c)
J2
oral
(d)
J2
oral
(e)
J2
oral
(f)
J2
oral
(g)
J2
oral
(h)
J2
DAPI
Elav
Msp130
oral

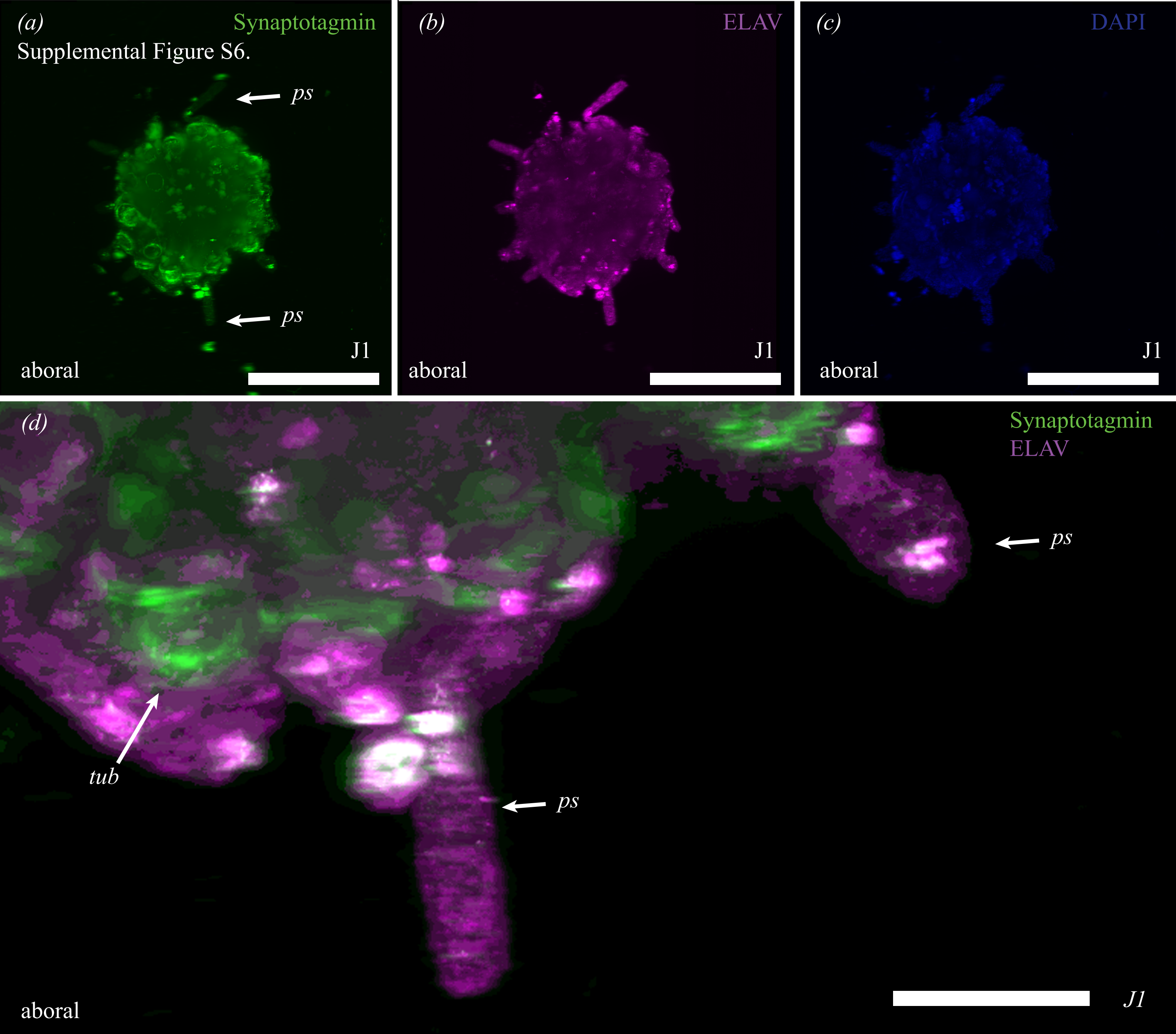
(a)
Synaptotagmin
Supplemental Figure S6.
ps
ps
J1
aboral
(b)
ELAV
J1
aboral
(c)
DAPI
J1
aboral
(d)
Synaptotagmin
ELAV
ps
tub
ps
aboral
J1

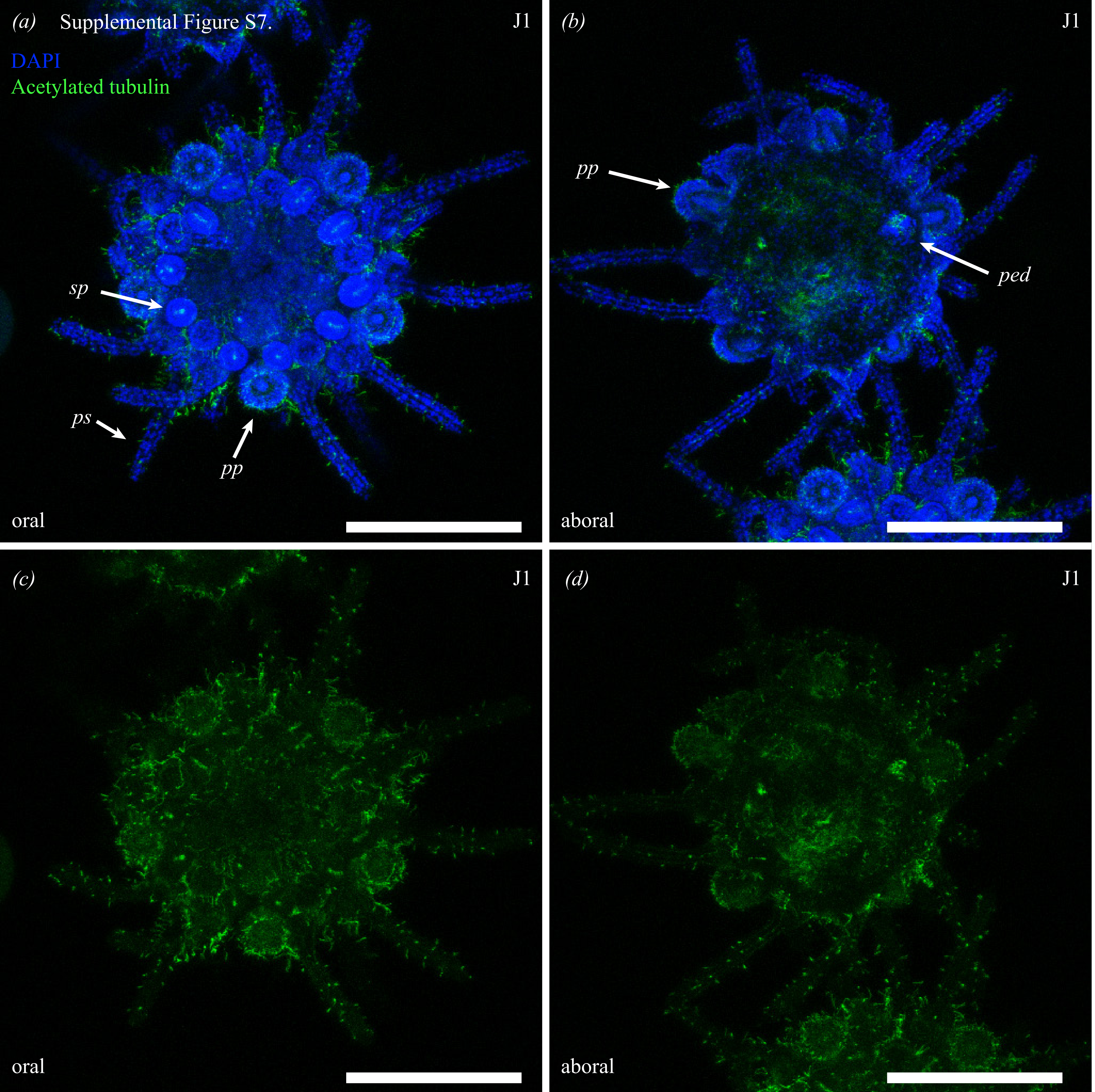
(a)
Supplemental Figure S7.
J1
DAPI
Acetylated tubulin
sp
ps
pp
oral
(b)
J1
pp
ped
aboral
(c)
J1
oral
(d)
J1
aboral

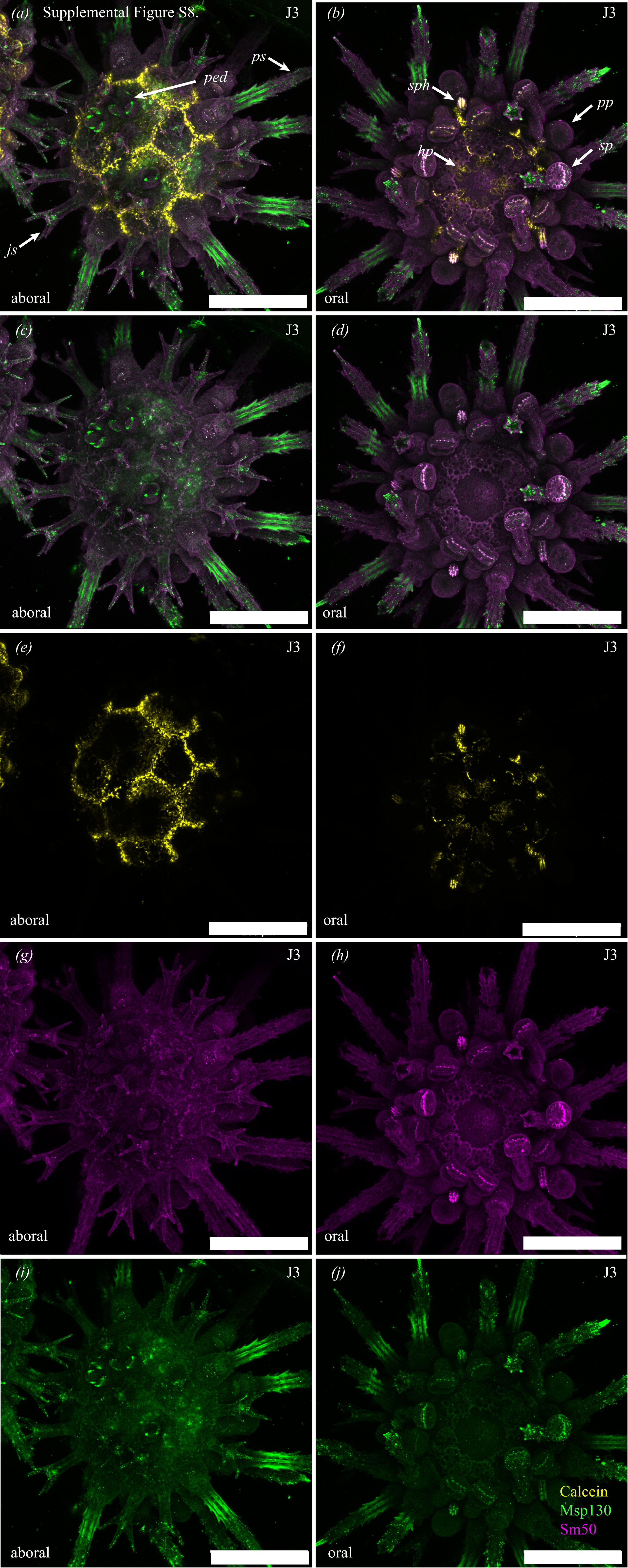
(a)
Supplemental Figure S8.
J3
ped
ps
js
aboral
(b)
J3
sph
pp
hp
sp
oral
(c)
J3
aboral
(d)
J3
oral
(e)
J3
aboral
(f)
J3
oral
(g)
J3
aboral
(h)
J3
oral
(i)
J3
aboral
(j)
J3
Calcein
Msp130
Sm50
oral

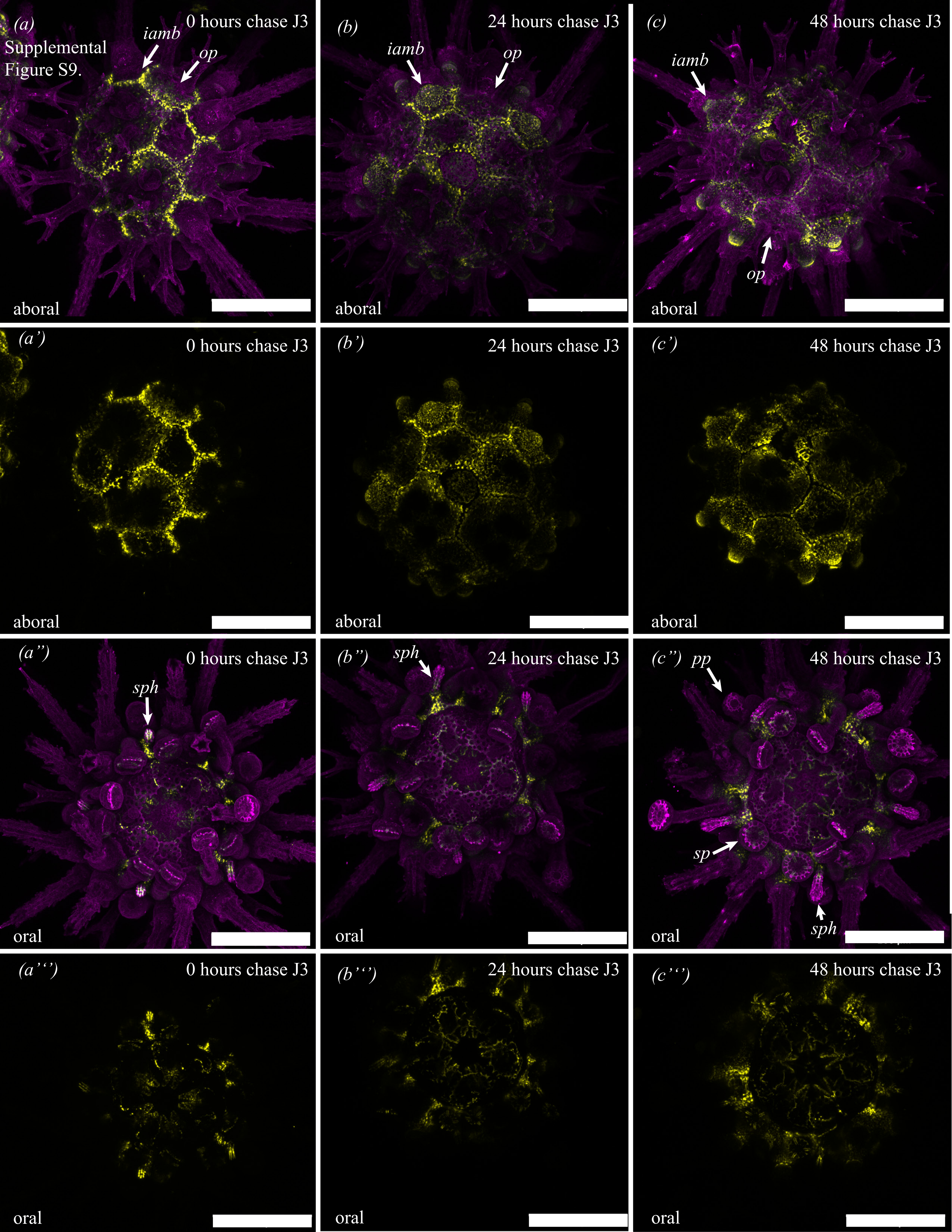

(a)
Supplemental
Figure S9.
0 hours chase J3
iamb
op
aboral
(b)
24 hours chase J3
iamb
op
aboral
(c)
48 hours chase J3
iamb
op
aboral
(a')
0 hours chase J3
aboral
(b')
24 hours chase J3
aboral
(c')
48 hours chase J3
aboral
(a")
0 hours chase J3
sph
oral
(b")
sph
24 hours chase J3
oral
(c")
pp
48 hours chase J3
sp
sph
oral
(a''')
0 hours chase J3
oral
(b''')
24 hours chase J3
oral
(c''')
48 hours chase J3
oral

Supplemental Figure S10.

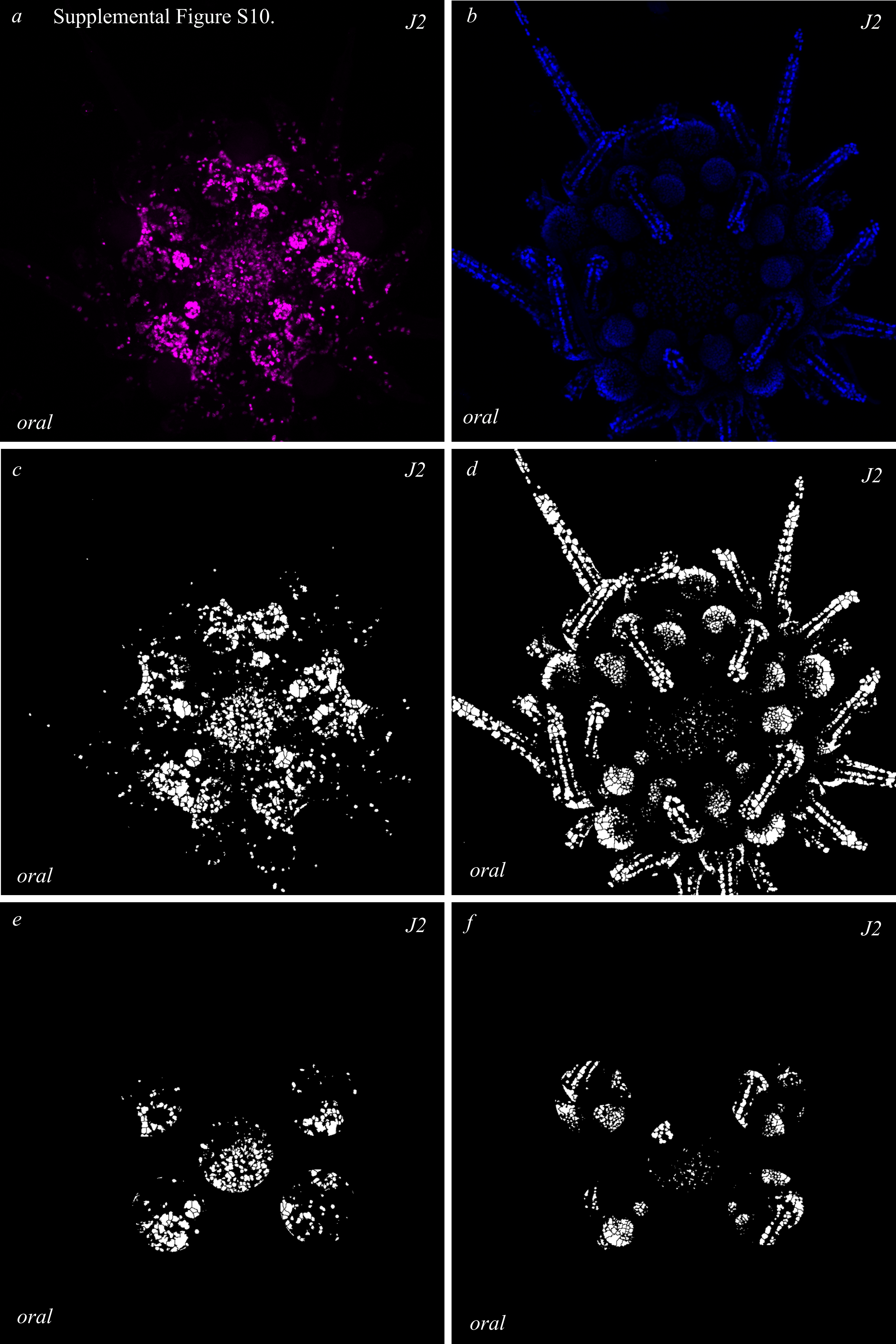

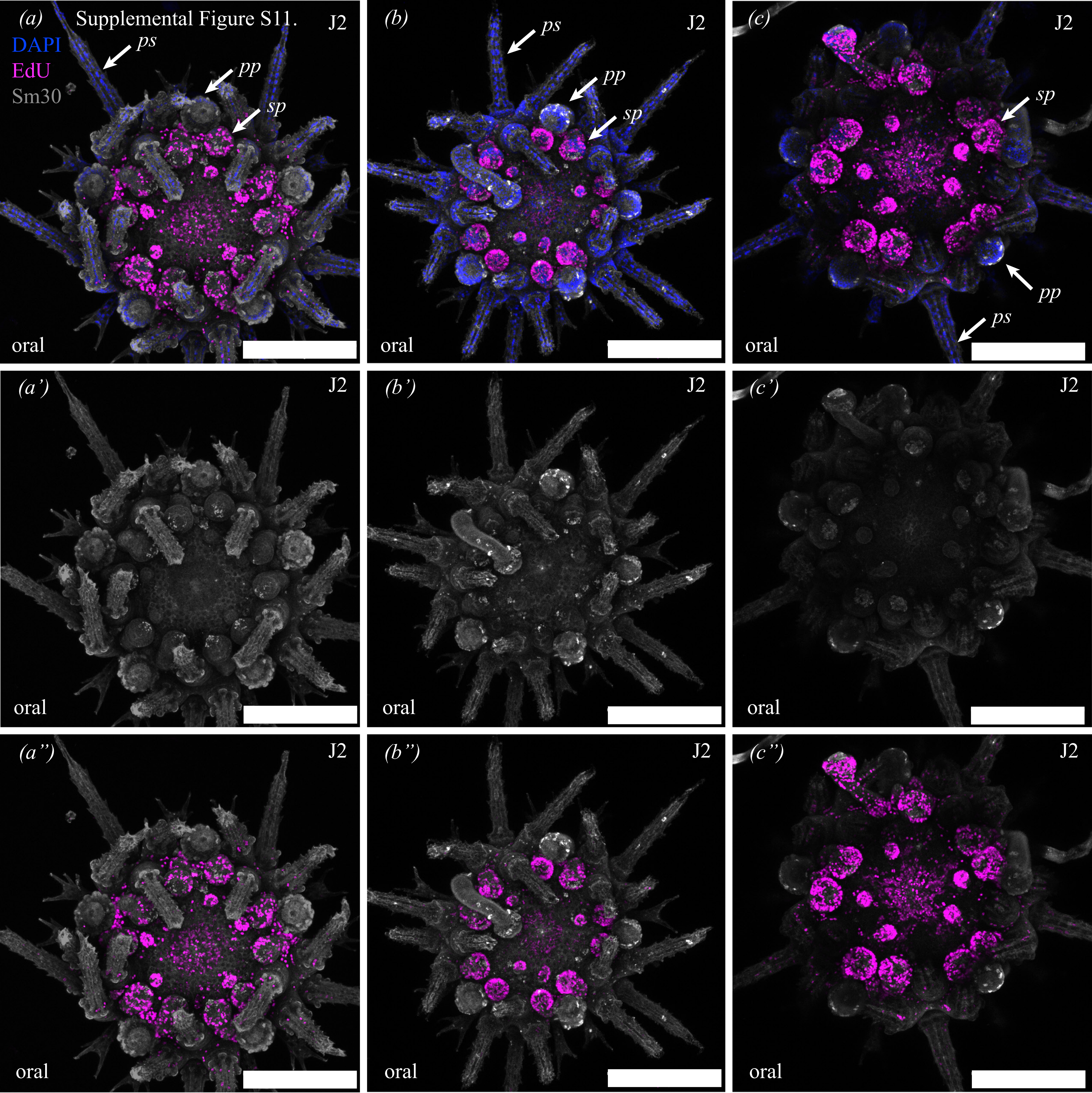
Supplemental Figure S11.
(a)
DAPI
EdU
Sm30
ps
pp
sp
J2
oral
(b)
(c)
(a')
(b')
(c')
(a")
(b")
(c")

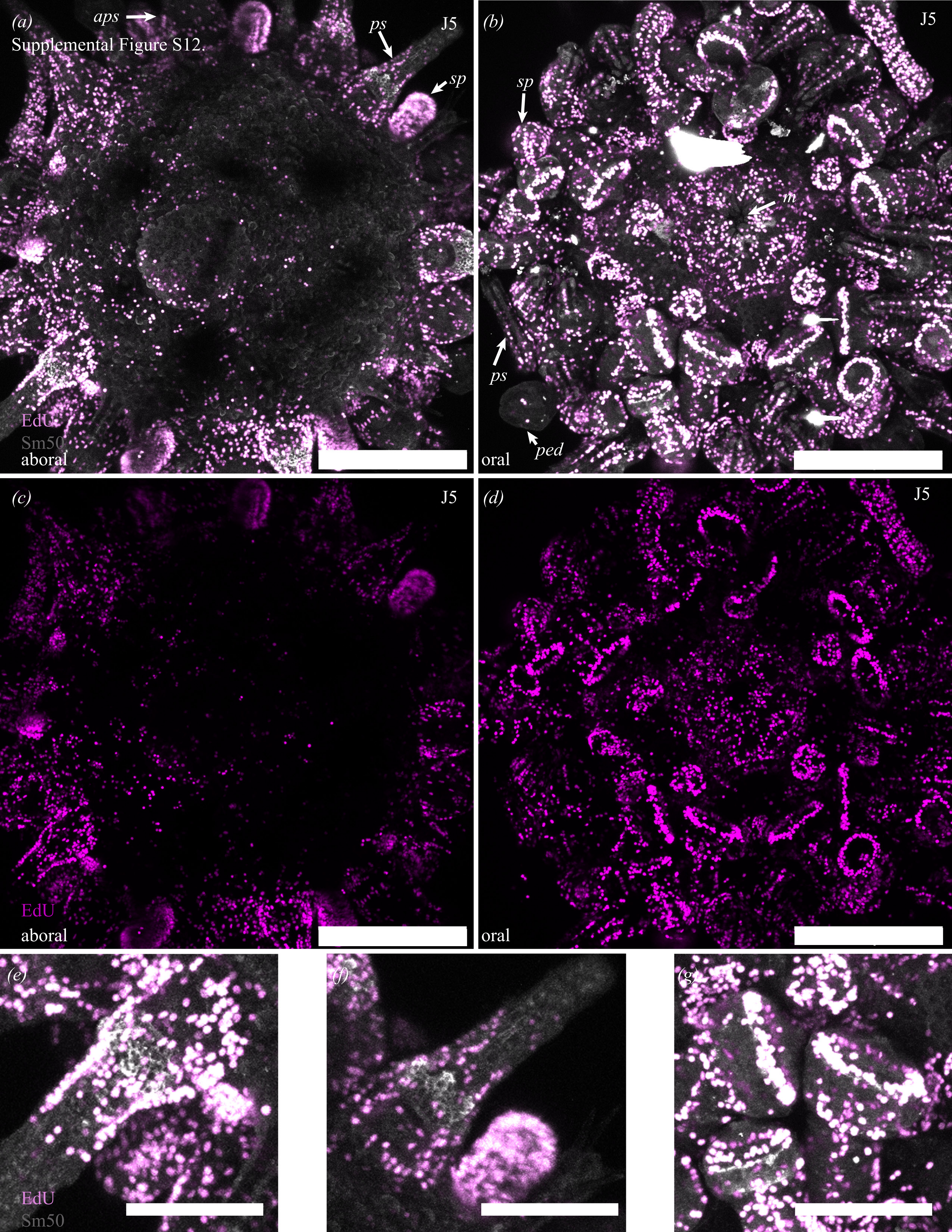

Supplemental Figure S12.
(a)
aps
ps
sp
J5
EdU
Sm50
aboral
(b)
J5
sp
m
ps
ped
oral
(c)
J5
EdU
aboral
(d)
J5
oral
(e)
EdU
Sm50
(f)
(g)

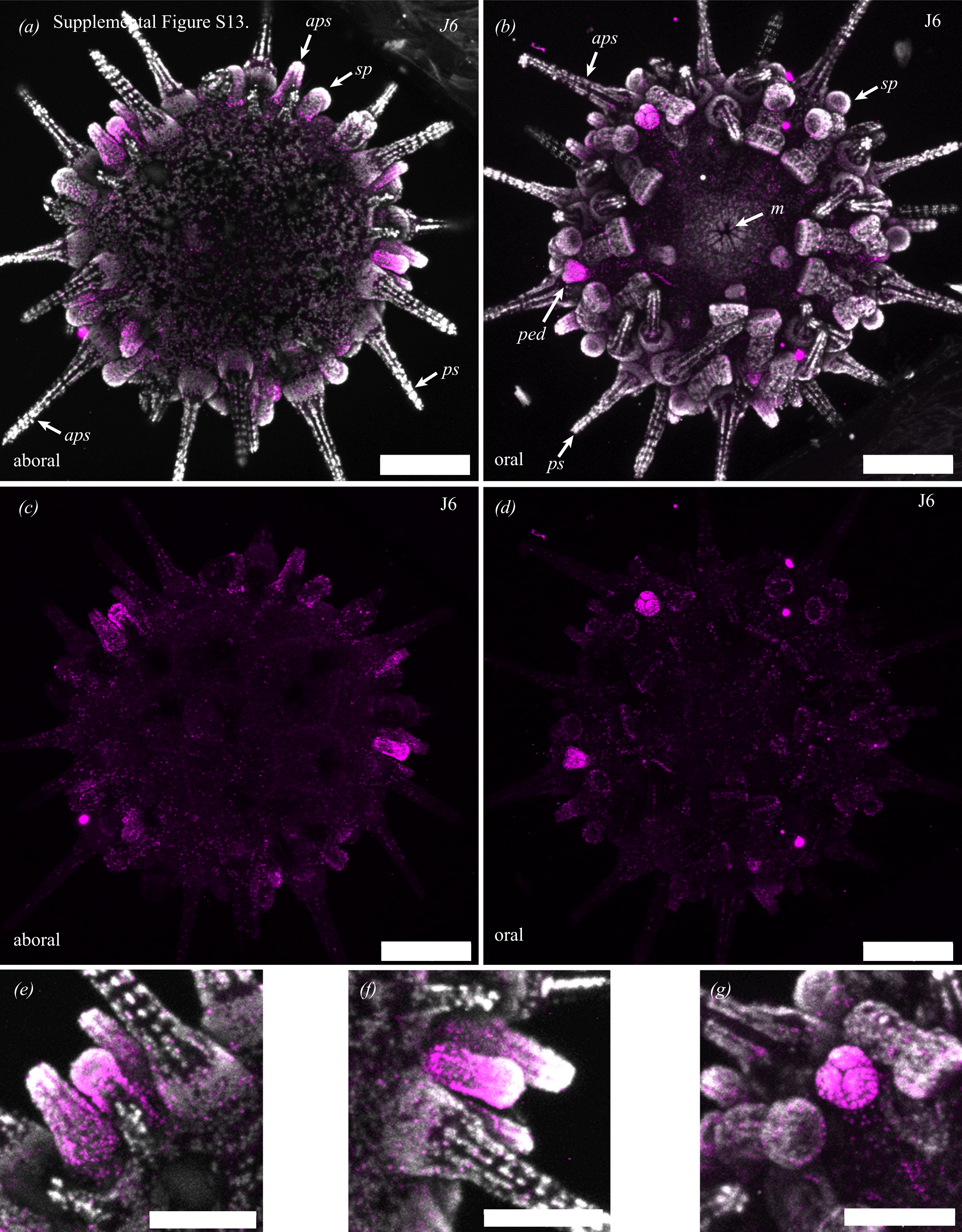
(a)
Supplemental Figure S13.
aps
sp
J6
ps
aps
aboral
(b)
J6
aps
sp
m
ped
ps
oral
(c)
J6
aboral
(d)
J6
oral
(e)
(f)
(g)
